# Parallel use of pluripotent human stem cell lung and heart models provide new insights for treatment of SARS-CoV-2

**DOI:** 10.1101/2022.09.20.508614

**Authors:** Rajeev Rudraraju, Matthew J Gartner, Jessica A. Neil, Elizabeth S. Stout, Joseph Chen, Elise J. Needham, Michael See, Charley Mackenzie-Kludas, Leo Yi Yang Lee, Mingyang Wang, Hayley Pointer, Kathy Karavendzas, Dad Abu-Bonsrah, Damien Drew, Yu Bo Yang Sun, Jia Ping Tan, Guizhi Sun, Abbas Salavaty, Natalie Charitakis, Hieu T. Nim, Peter D Currie, Wai-Hong Tham, Enzo Porrello, Jose Polo, Sean J. Humphrey, Mirana Ramialison, David A. Elliott, Kanta Subbarao

## Abstract

SARS-CoV-2 primarily infects the respiratory tract, but pulmonary and cardiac complications occur in severe COVID-19. To elucidate molecular mechanisms in the lung and heart, we conducted paired experiments in human stem cell-derived lung alveolar type II (AT2) epithelial cell and cardiac cultures infected with SARS-CoV-2. With CRISPR- Cas9 mediated knock-out of ACE2, we demonstrated that angiotensin converting enzyme 2 (ACE2) was essential for SARS-CoV-2 infection of both cell types but further processing in lung cells required TMPRSS2 while cardiac cells required the endosomal pathway. Host responses were significantly different; transcriptome profiling and phosphoproteomics responses depended strongly on the cell type. We identified several antiviral compounds with distinct antiviral and toxicity profiles in lung AT2 and cardiac cells, highlighting the importance of using several relevant cell types for evaluation of antiviral drugs. Our data provide new insights into rational drug combinations for effective treatment of a virus that affects multiple organ systems.

**One-sentence summary:** Rational treatment strategies for SARS-CoV-2 derived from human PSC models

## Introduction

Coronavirus disease 19 (COVID-19) is primarily a respiratory disease. About 80% of infections are clinically mild or asymptomatic. Progression to severe illness is associated with lower respiratory tract involvement with pneumonia and acute respiratory distress syndrome (1, 2). Pulmonary fibrosis has been reported in survivors of COVID-19 (3).

In addition to pulmonary disease, cardiovascular, renal, digestive and neurological complications are reported (4). Cardiac complications include arrhythmias, thromboembolism, acute myocardial injury associated with elevated levels of cardiac troponin and electrocardiographic abnormalities (5, 6). Magnetic resonance imaging has shown cardiac involvement in up to 78% of recovered COVID-19 patients, and ongoing myocardial inflammation in 60% of patients (7, 8). Moreover, a meta-analysis of studies in COVID-19 patients found that myocardial injury was significantly associated with increased mortality (9). While cardiac damage during COVID-19 is predominantly thought to be due to an over-exuberant immune response, studies of autopsy tissue from patients who died from COVID-19 have detected viral RNA and Spike (S) antigen in the heart (10–12).

SARS-CoV-2 mediates infection by binding of the S protein to its receptor, angiotensin converting enzyme 2 (ACE2) on the host cell (13). The S protein is cleaved into two domains, S1 and S2 by host cell proteases, including furin (14). Following attachment, fusion with the cell membrane requires further proteolytic cleavage at the S2’ site to activate the fusion peptide (15). This is mediated extracellularly by serine proteases including Transmembrane protease, serine 2 (TMPRSS2) or in endosomes by Cathepsin L (15–17). Thus, ACE2 is a critical determinant of the tissue tropism of SARS-CoV-2, as are the presence of proteases and/or endosomal pathways for activation of the fusion activity of the S protein. ACE2 is expressed in many human tissues including the lungs, nose, cornea, heart, kidney, esophagus, gastrointestinal tract, liver, gallbladder placenta and testis, with high ACE2 expression observed in the nasal epithelium, lungs, ileum and heart (18–23). Tissue sites containing cells that co-express ACE2 and TMPRSS2 include the nose, lungs, kidney, gastrointestinal tract and the gallbladder (18, 22), while cells co- expressing ACE2 and Cathepsin L are found within the lung, heart and the gastrointestinal tract (22). Following entry, the virus interacts with the cellular machinery to complete its replication cycle and triggers a host cell response that can vary in different organs.

We sought to elucidate the molecular mechanisms of SARS-CoV-2 infection in lung and heart using human stem cell-derived lung and cardiac cells. Human pluripotent stem cells (hPSCs) including both human embryonic stem cells (hESCs) and induced pluripotent stem cells (hiPSCs) have been used to generate functional human cells, tissue and organoids to model human disease. We and others have generated stem cell-derived lung alveolar type II (AT2) epithelial cell and cardiac cultures that can be productively infected with SARS-CoV-2 (17, 24–26). We hypothesized that paired experiments in SARS-CoV-2 infected lung and cardiac cells would reveal important similarities and differences in viral and host factors that could inform treatment of COVID-19 and its complications. We used CRISPR-Cas9 mediated knock-out of ACE2 and demonstrated that ACE2 was essential for SARS-CoV-2 infection of both cell types. Small molecule inhibitors revealed distinct mechanisms of SARS-CoV-2 entry. We identified differential cellular responses to SARS-CoV-2 infection by transcriptome profiling and phosphoproteomics and further demonstrated the utility of these stem cell-derived models for screening antiviral compounds for anti-SARS-CoV-2 activity. Our findings provide new insights into treatment strategies for COVID-19.

## Results

### SARS-CoV-2 productively infects human stem cell-derived lung AT2 and cardiac cultures

hESC and iPSC-derived cardiomyocyte (27, 28) and AT2 lung (29) cultures were generated to develop *in vitro* models of SARS-CoV-2 infection (Figure 1A). Lung AT2 cultures expressed high levels of lung development homeobox protein NKX2-1 and low levels of the type I alveolar cell marker Aquaporin-5 (AQP5) confirming the presence of low numbers of alveolar type I cells (Figure S1A). Gene expression profiling of the lung AT2 and cardiac cells demonstrated transcriptional profiles consistent with high levels of AT2 cells and cardiomyocytes, respectively (Figure S1B, C). For example, key cardiac genes such as those encoding myofilament proteins (*TNNT2*, *MYH7*, *MYL2* and *TTN*) and cardiomyocyte transcription factors (*HAND2*, *MEF2C* and *NKX2-5*) were expressed in cardiac cultures whilst the lung AT2 cultures expressed lung-specific development markers (*NKX2-1*, *SOX9* and *GATA6*) and signaling genes (*SHH* and *BMP4*).

**Figure 1.**
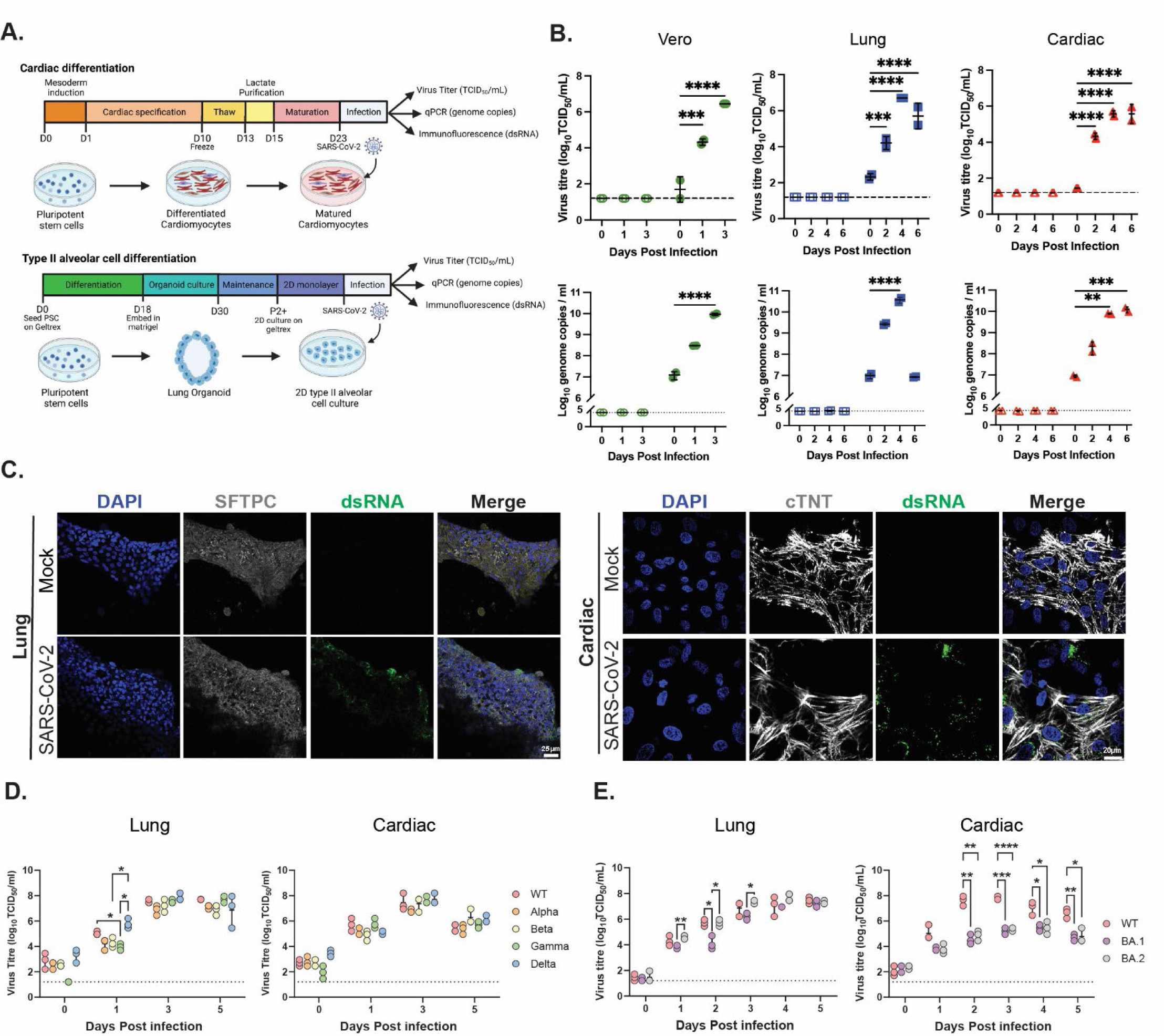
SARS-CoV-2 infection of cardiac and lung AT2 cells is mediated by ACE2. (A) Schematic of cell differentiation and infection protocols. (B) Viral titres and genome copies in SARS-CoV-2 (VIC01) infected Vero, lung AT2 (H9) and cardiac (NKX2-5) culture supernatants. (C) Representative fluorescent confocal microscopy images of dsRNA (green) expression in VIC01 infected lung AT2 (SFTPC-positive) and cardiac (cTNT-positive) cells at 3 dpi. Titres in supernatants from cardiac and lung AT2 cells infected with VIC01 (WT) compared to cells infected with Alpha, Beta, Gamma and Delta variants (D) or Omicron (BA.1 and BA.2) variants (E).

To determine susceptibility to SARS-CoV-2 infection, lung AT2 and cardiac cells were inoculated with the ancestral strain of SARS-CoV-2. Vero cells were included in the experiment as a positive control for virus infection. As expected, SARS-CoV-2 productively infected Vero cells with virus titres and E gene copies, determined by infectivity assay and qPCR respectively, peaking at 3 days post-infection (dpi) (Figure 1B). SARS-CoV-2 showed robust virus replication in lung AT2 cells, with virus titres and E gene copies, peaking at 4 dpi (Figure 1B). Immunostaining for double stranded RNA (dsRNA) showed evidence of SARS-CoV-2 replication at 3 dpi (Figure 1C). Cardiac cell cultures differentiated from hESCs (*NKX2-5^eGPF/w^* (30)) were also susceptible to SARS-CoV-2 infection with virus titres and E gene copies peaking slightly later than AT2 cells between 4 and 6 dpi (Figure 1B). Cardiac cells generated from *ALPK3* knockout hESCs that model hypertrophic cardiomyopathy (31), were similarly susceptible to SARS-CoV-2 infection with a peak in virus titres at 4 dpi (Figure S1D). Immunostaining showed SARS-CoV-2 dsRNA in both cardiomyocyte and non-cardiomyocyte cells within the cardiac monolayer cultures (Figure 1C). Furthermore, cardiac cultures infected with SARS-CoV- 2 stopped contracting at 4 dpi (Video S1). Overall, these data show that SARS-CoV-2 replicates efficiently in cardiac and lung AT2 cells derived *in vitro* from hPSCs, consistent with published reports (17, 24).

To determine whether cardiac and lung AT2 cells are susceptible to infection by SARS-CoV-2 variants, virus titres and E gene copies were assessed following infection with Alpha, Beta, Gamma, Delta and Omicron (BA.1 and BA.2) variants. Although all variants were able to infect the lung AT2 cells, a modest difference was observed following Delta infection in lung AT2 cells at 1 and 3 dpi (Figure 1D, S1E) but titres were comparable to all other variants by 5 dpi. In cardiac cells, a slightly lower virus titer was observed following infection with the Alpha variant at 3 dpi (Figure 1D). However, this was not observed for E gene copies where WT levels were higher than other variants on 1 and 3 dpi (Figure S1E). In lung AT2 cells, the Omicron subvariants showed similar titres to the WT virus, although BA.1 showed a modest reduction in virus titres and E gene copies compared to BA.2 (Figure 1E, S1F). In cardiac cells, the BA.1 and BA.2 variants showed significantly lower virus titres compared to the WT virus at 2-5dpi (Figure 1E). However, this difference in replication was not observed for E gene copies (Figure S1F). These data confirm that hPSCs derived *in vitro* models can be used to study all variants and that SARS-CoV-2 variants infect cardiac and lung cells efficiently.

### SARS-CoV-2 infection in lung AT2 and cardiac cells is dependent on ACE2

ACE2 is the functional receptor for sarbecoviruses, the SARS coronavirus family. To confirm that ACE2 is required for SARS-CoV-2 infection in cardiac and lung AT2 cells, we generated two *ACE2* knockout (KO) hPSC lines (H9 and MCRIi010-A) engineered via CRISPR/Cas9 to remove the first coding exon of ACE2 (Figure S2A and B). ACE2 protein and *ACE2* transcript expression was not detected in either lung AT2 or cardiac cultures differentiated from the *ACE2* knockout line (Figure S2C and D). Flow cytometric analysis confirmed that the genetically modified cells maintained their expression of pluripotency markers and showed similar differentiation capacity (Figure S2E and F). Furthermore, immunofluorescence staining in ACE2 KO cardiac or lung AT2 cells showed that ACE2 expression was undetectable (Figure S2H).

Following confirmation that ACE2 was knocked out in H9 and MCRIi010-A lines, cardiomyocyte and AT2 differentiation protocols were performed. The ACE2 KO cardiac or lung AT2 cells could not be productively infected with SARS-CoV-2, as shown by the inability to recover infectious virus and absence of dsRNA staining (Figure 2A, S2G-S2H). For independent confirmation of the role of ACE2, we used a combination of two previously described α-ACE2 antibodies at doses between 2-40 µg/mL (23). In cardiac cultures, 2 μg/mL of α-ACE2 antibodies was sufficient to completely block SARS-CoV-2 infection (Figure 2B). In contrast, treatment with the α-ACE2 antibody cocktail blocked infection in a concentration-dependent fashion in lung AT2 cells (Figure 2B). Overall, our data demonstrate that SARS-CoV-2 infection in lung AT2 and cardiac cells is dependent on ACE2 for virus entry.

**Figure 2.**
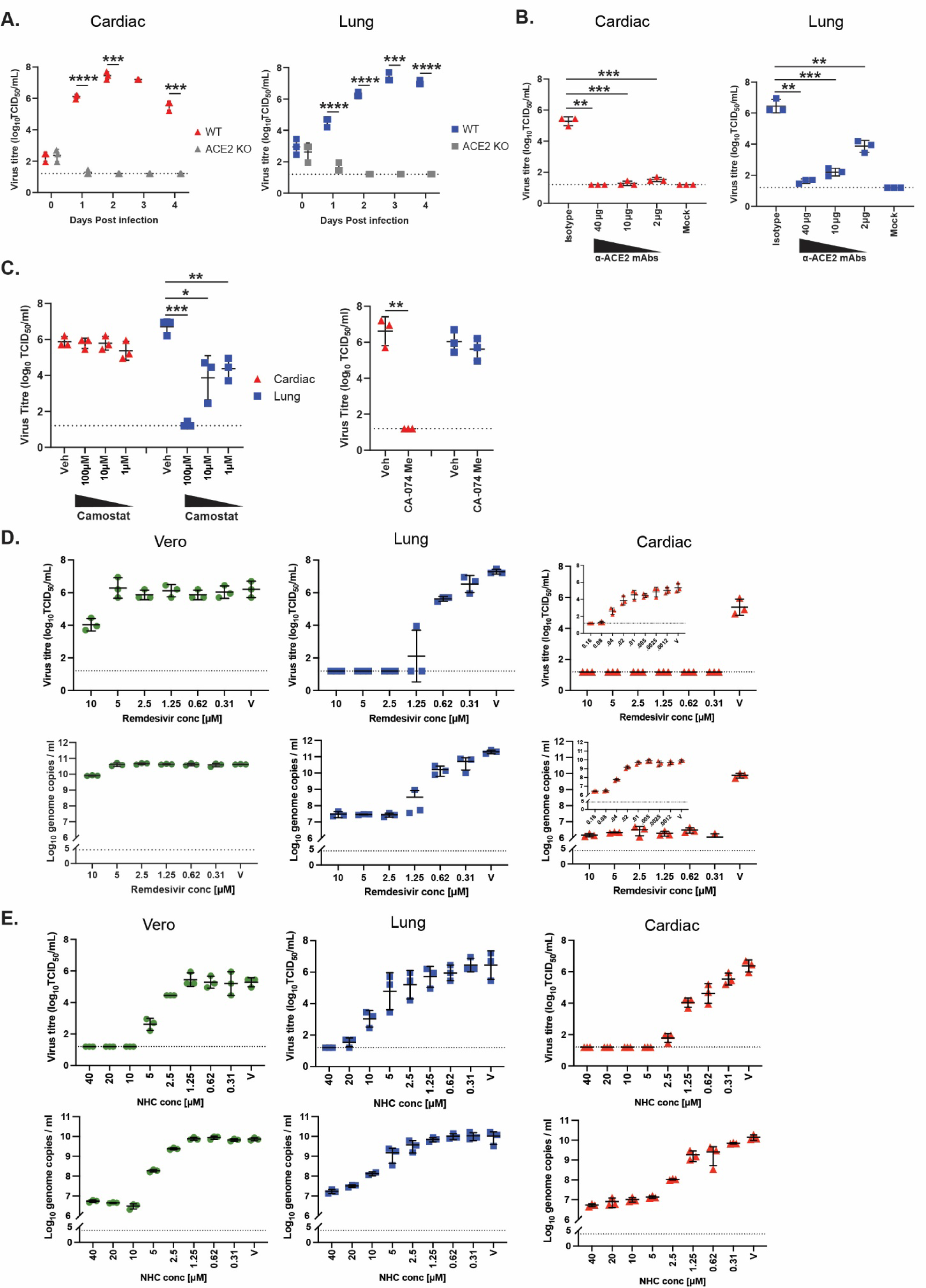
SARS-CoV-2 entry and antiviral sensitivity is different between lung AT2 and cardiac cells. (A) Virus titer in supernatants from SARS-CoV-2 (VIC01) infected H9- derived WT and *ACE2* KO cardiac and lung AT2 cells. (B) Virus titer at 3 dpi in supernatants from lung AT2 (H9) and cardiac cells (NKX2-5) treated with two α-ACE2 antibodies or a human IgG1 isotype control before infection with SARS-CoV-2. (C) Virus titer at 3 dpi in supernatant from cardiac cells (red triangles) and lung AT2 cells (blue squares) infected with SARS-CoV-2 (VIC01) in the presence of Camostat, CA-074 or DMSO (vehicle control). Virus titres and genome copies at 3 dpi in supernatant from Vero, lung AT2 and cardiac cells infected with SARS-CoV-2 VIC01 in the presence of various concentrations of Remdesivir (D) or NHC (E).

### SARS-CoV-2 utilizes differential entry mechanisms in lung AT2 and cardiac cells

Following attachment to the host cell receptor, coronaviruses require proteolytic activation of the spike protein to mediate fusion of the viral and host cell membranes. SARS-CoV-2 can be activated for fusion at the cell membrane through transmembrane protease, serine 2 (TMPRSS2) mediated cleavage of the (S2’) site or through endocytosis, where endosomal acidification triggers Cathepsin L (CSTL)-activated fusion between the viral and endolysosomal membranes. While lung AT2 cells had high expression of *TMPRSS2*, cardiac cells did not express *TMPRSS2* (Figure S3A, B). In contrast, both lung AT2 and cardiac cells expressed *CSTL* at high levels (Figure S3A, B) suggesting that SARS-CoV-2 may enter cardiac cells via the endocytosis pathway.

To determine the mechanism of entry, cells were infected in the presence of the TMPRSS2 inhibitor Camostat mesylate or the Cathepsin B/L inhibitor CA-074 Me. Camostat treatment (100, 10 and 1 μM) led to a significant reduction in virus titer and genome copies in lung AT2 but not in cardiac cells (Figure 2C and S3C) showing that infection in lung AT2 cells requires TMPRSS2 cleavage. In contrast, 25 μM of CA-074 Me led to a significant reduction in virus titer and genome copies in cardiac but not in lung AT2 cells (Figure 2C and S3C), confirming that the viral entry in cardiac cells requires the endosomal pathway. Thus, consistent with the gene expression data, SARS-CoV-2 utilizes distinct entry pathways in lung AT2 and cardiac cells.

### Antiviral compounds show differential activity in lung and cardiac cells compared to traditional cell lines

To determine whether SARS-CoV-2 antiviral drugs show similar activity in lung AT2 and cardiac cells, we investigated two sets of small molecules: (a) drugs that are approved for treatment of COVID-19 including Remdesivir and NHC (β-D-N4- hydroxycytidine), the prodrug of Molnupiravir and (b) drugs that were reported to show antiviral activity *in vitro* but are still under clinical investigation or were not found to be effective in clinical trials including favipiravir, tizoxanide (the active form of the antiparasitic agent nitazoxanide), and chloroquine.

Remdesivir and NHC significantly inhibited viral replication in both cardiac and lung AT2 cells (Figure 2D, E). In Vero cells, remdesivir inhibited virus replication at a concentration of 10 μM while in lung AT2 cells, the drug completely inhibited virus replication at concentrations as low as 2.5 µM (IC_50_ = 0.55 µM) and antiviral activity in cardiac cells was far more potent, with complete inhibition of viral replication at 0.08 μM (IC_50_=0.016 µM) (Figure 2D). Similarly, NHC was more effective in inhibiting virus replication in cardiac cells compared to lung AT2 and Vero cells, with complete virus inhibition observed at concentrations as low as 5 µM (IC_50_=0.42 µM) (Figure 2E) while viral replication was inhibited in Vero and lung AT2 cells with similar dose-response kinetics (IC_50_=2.4 µM and 3.54 µM, respectively) (Figure 2E). In terms of toxicity, reduced viability of both lung AT2 and cardiac cells was observed with remdesivir concentrations above 5 µM but not in Vero cells (Figure S4A). NHC was noticeably less toxic than remdesivir in all three cell types, with toxicity observed at the concentration of 40 μM in cardiac and lung AT2 cells (Figure S4B). These results highlight the variability in both antiviral efficacy and cytotoxicity in different cell types and emphasize the relevance of using human stem-cell derived models or human cells over Vero cells for assessment of antiviral drugs for COVID-19.

Favipiravir, piperaquine and tizoxanide showed no antiviral activity against SARS-CoV-2 in cardiac, lung AT2 and Vero cells up to a concentration of 10 µM (Figure S4C). Of these three compounds, tizoxanide was toxic in Vero cells but not in cardiac or lung AT2 cells (Figure S4C). Chloroquine was not toxic in any of the cell types and inhibited SARS-CoV-2 virus replication in cardiac cells (at 10 µM) but not in lung AT2 or Vero cells (Figure S4C), consistent with our observation that SARS-CoV-2 entry in cardiac cells utilizes the endosomal pathway. Chloroquine also inhibited virus replication in the *ALPK3* KO cardiomyopathy model (Figure S4C). Together, these data indicate that the activity of antiviral drugs differs between lung AT2 and cardiac cells compared to conventionally used Vero cells.

### SARS-CoV-2 induces different transcriptional responses in lung AT2 and cardiac cells

Based on our observation of distinct entry pathways and antiviral activity between lung AT2 and cardiac cells, we hypothesized that they would show diverse responses to SARS-CoV-2 infection. RNA-sequencing (RNA-seq) was performed on cell lysates from hESC-derived lung AT2 and cardiac cells infected with SARS-CoV-2 at 1 and 3 dpi. A principal component analysis (PCA) plot separated the tissue types and demonstrated a progressive separation between mock-infected and virus-infected cells particularly by 3 dpi (Figure 3A). At 1 dpi, no differentially expressed genes (DEGs) were observed in lung AT2 cells, and only 5 in cardiac cells (data not shown). However, at 3 dpi, 1154 and 992 DEGs were obtained for cardiac and lung AT2 cells, respectively, with specific lung and cardiac specific DEGs. As shown in Figure 3B, of the DEGs between the 2 cell-types, 202 were overlapping (intersect), 952 genes were differentially expressed in the cardiac cells only (cardiac unique) and 790 genes were differentially expressed in the lung AT2 cells only (lung unique). Additionally, we found that SARS-CoV-2 infection did not change the expression profile of lung development genes in lung AT2 cells and cardiac development genes in cardiac cells at 1 and 3 dpi (Figure S1B and C).

**Figure 3.**
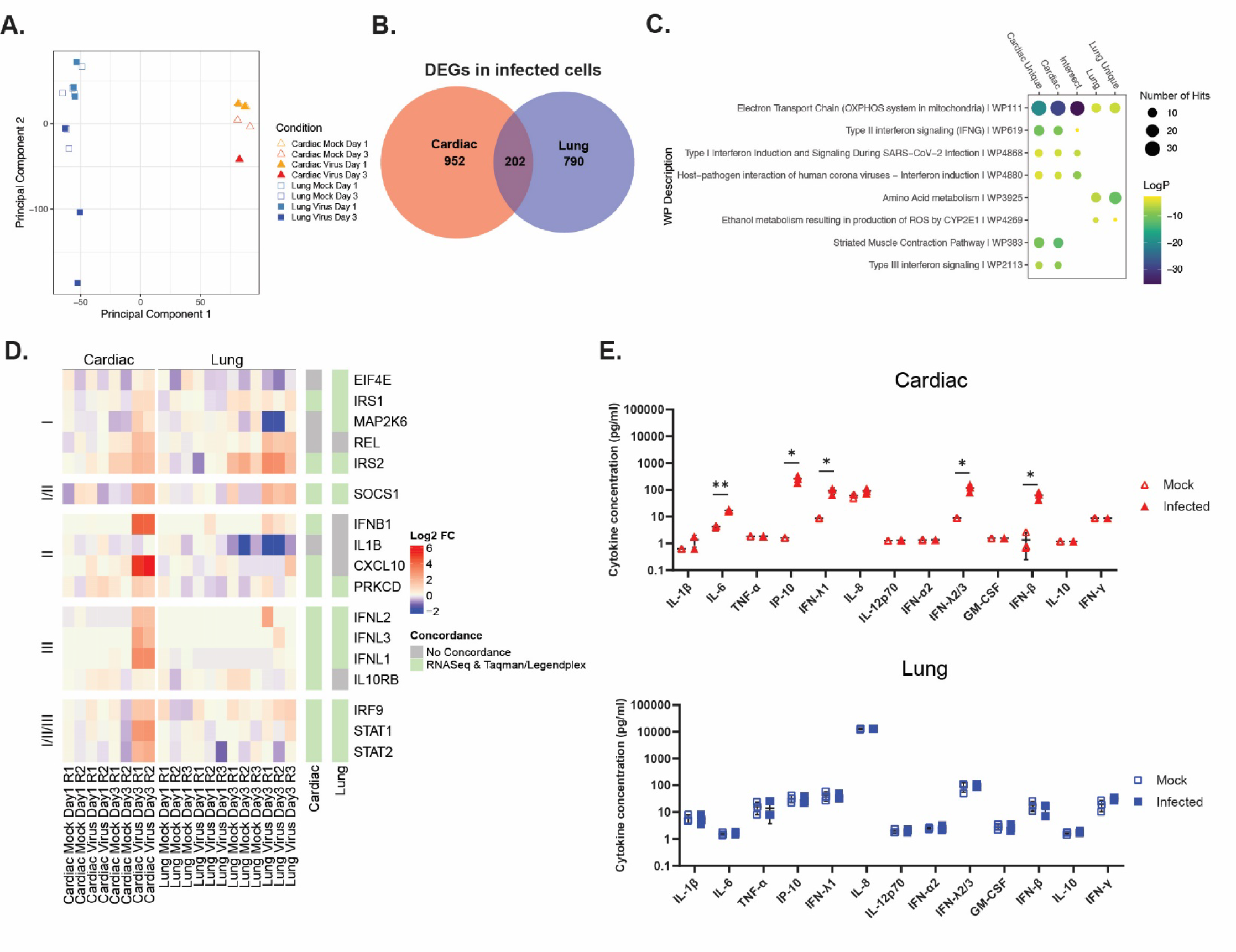
Cardiac, but not lung AT2 cells, have a robust interferon signature following SARS-CoV-2 infection. (A) Principal Component Analysis (PCA) plot of uninfected (mock) and SARS-CoV-2 infected (virus) cardiac and lung AT2 cells at 1 and 3 dpi. (B) Venn diagram depicting the number of overlapping and unique differentially expressed genes compared to mock-infected. (C) Top representative WikiPathway (WP) terms of the enriched genes. Circle size is proportional to the number of genes that matched the pathway and color represents the LogP value as calculated by Metascape. (D) Heat map in Log2 fold change (FC) of representative interferon genes (compared to representative mock-infected samples at 1 dpi) separated by type as defined by WikiPathways. Columns on the right show the concordance between direction of fold change of RNAseq data and the LegendPlex/Taqman assays. For 1 dpi and 3 dpi, 3 lung samples and 2 cardiac samples were analyzed per group (mock, virus). (E) Cytokine concentrations in the supernatants of mock and SARS-CoV-2 infected cardiac and lung AT2 cells at 3 dpi.

WikiPathway enrichment analysis of DEG subsets identified that the biological processes impacted by SARS-CoV-2 infection in both lung AT2 and cardiac cells (i.e. intersect subset), included Electron Transport Chain (OXPHOS system in mitochondria) (WP111), Host-pathogen interaction of human coronaviruses – Interferon (IFN) induction (WP4880) and Type I and II IFN signaling pathways (Figure 3C; Figure S5A, B). However, only 3 genes in the “Intersect” subset involved Type II IFN signaling, indicating only a relatively small response. In contrast, the “cardiac unique” gene subset, included an overall upregulation of genes in Type III interferon signaling (WP2113) and downregulation of genes in the Striated Muscle Contraction Pathway (WP383), consistent with the cessation of contractility of SARS-CoV-2 infected cultures (Video S1). In the “lung unique” subset, we observed an overall downregulation of genes in Amino Acid metabolism (WP3925) and upregulation of genes in Ethanol metabolism resulting in production of reactive oxygen species (ROS) by CYP2E1 (WP4269) (Figure 3C).

Key differences in the transcriptional responses to SARS-CoV-2 infection between lung AT2 and cardiac cells were the IFN pathways that were activated (Figure 3C, D). By 3 dpi, genes from the Type I, II and III IFN pathways (*IFNB1*, *CXCL10*, *PRKCD*, *IFNL1*, *IFNL2* and *IFNL3*) were upregulated in cardiac but not lung AT2 cells. Concordant with the observed increase in transcription by RNASeq, cytokine analysis showed a significant induction of IL-6, IP-10, IFN-λ1, IFN-λ2/3 and IFN-β in infected cardiac cells, but not lung AT2 cells, at 3 dpi compared to mock-infected controls (Figure 3E). Furthermore, qPCR showed induction of *IFITM3*, *IFN-β*, *STAT1* and *STAT2* in infected cardiac cells, and not lung AT2 cells, compared to mock-infected cells at 3 dpi (Figure S5C). Network analysis of the GO term response to interferon gamma showed multiple genes within the network were upregulated in infected cardiac but not lung AT2 cells (Figure S5D). Taken together, SARS-CoV-2 infection in cardiac induces robust Type I, II and III IFN responses but not in lung AT2 cells.

### SARS-CoV-2 activates druggable kinases in lung AT2 and cardiac cells

We measured the phosphoproteome to determine the signaling responses of lung AT2 and cardiac cells to SARS-CoV-2 infection. Cells were either mock-infected or infected with SARS-CoV-2 and sampled at 0-, 18-, 24- and 72- hours post inoculation (Figure 4A). Only viral phosphorylation sites were analyzed at the 72 hours timepoint. By employing recent advances in phosphoproteomics technologies (32), we quantified >32,000 phosphopeptides in at least three samples (Figure 4B). The phosphoproteomes clustered primarily by tissue origin in PCA (PC1), emphasizing the cell-type specific nature of signaling (Figure 4C). Time since infection was the second largest factor affecting clustering by PCA (on PC2), due to the long time-points studied. For this reason, a ‘mock’ condition was sampled at every timepoint, enabling identification of infection-driven signaling. SARS-CoV-2 infection extensively regulated cellular signaling, with at least 250 phosphopeptides regulated at each timepoint (Figure 4D). Regulation of the phosphoproteome was most extensive in cardiac cells at 18 hours post-infection (>900 phosphopeptides altered). Remarkably only 12 and 15 phosphopeptides were commonly regulated between the cardiac and lung AT2 cells at the 18- and 24- hour timepoints, respectively (Figure 4E). This low overlap of regulation was also observed at the transcriptional level and indicates that the response to SARS-CoV-2 is highly contextual and depends strongly on the cell type or tissue of origin.

**Figure 4.**
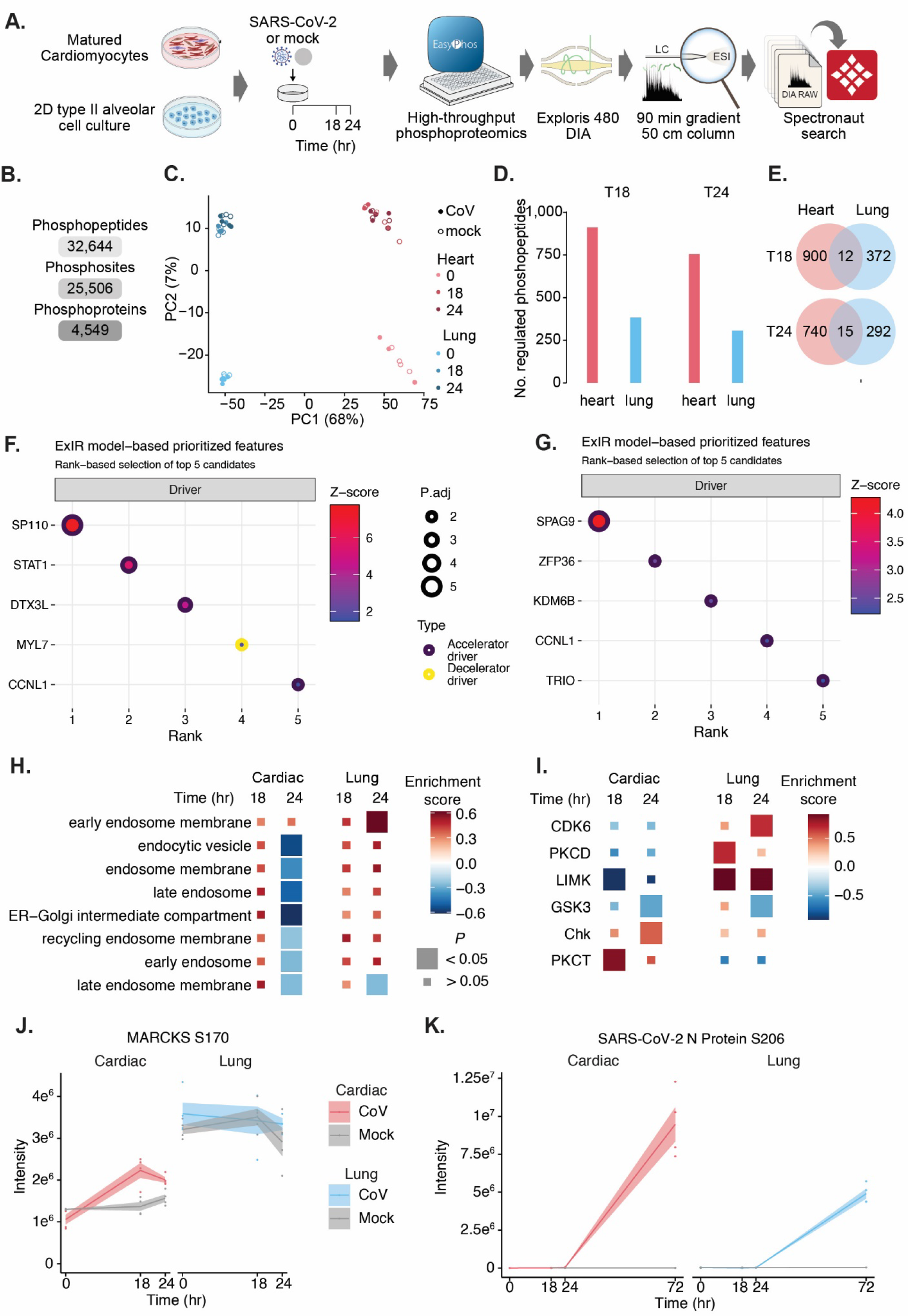
SARS-CoV2 infection alters different signaling networks in heart and lung organoids. (A) Schematic of phosphoproteomics workflow. (B) Number of phosphorylated peptides, sites and proteins found in at 3 samples. (C) PCA plot of median-normalized phosphoproteome replicates from uninfected (mock) and SARS-CoV- 2 infected (CoV) cardiac and lung AT2 cells at 18 and 24 hours post infection. Number of significantly (Padj < 0.05, FC >1.5) regulated phosphopeptides per timepoint (D) and their overlap (E) between cell types. Top 5 predicted drivers following SARS-CoV-2 infection in lung and cardiac cells (F) and in lung cells (G) at 72h post-infection. Adjusted P-values (P.adj) are calculated based on the computation of Z-score probability distributions of a molecule to be a driver and adjusted using the Benjamini and Hochberg algorithm. (H) GO cellular components enriched in at least one condition (P < 0.05) that are related to endosomes. (I) Kinases with enriched substrates, which were selected for targeting. (J) Abundance of the PKC substrate MARCKS S170. (K) Abundance of the SRPK1 substrate SARS-CoV2 nucleocapsid protein S206.

To corroborate this, we sought to identify the molecular drivers of the SARS-CoV- 2 infection response, by integrating RNA-seq and phosphoproteomics datasets using ExIR (https://influential.erc.monash.edu/). This analysis revealed that the top 3 drivers of SARS-CoV-2 infection response in cardiac cells (*SP110*, *STAT1* and *DTX3L*) were activators of the interferon response (Fig. 4F, G). In contrast, no strong interferon pathway activators were identified in the top drivers of the infection response in lung AT2 cells, confirming the specificity of the interferon response only in cardiac cells. To determine the cellular context of SARS-CoV2-regulated signaling, we tested for enrichment of cellular components (Figure S6A). Endosomal components almost uniquely enriched in the SARS-CoV2 response of cardiac cells, and not lung cells (Figure 4H, Figure S6A). This mirrors the pattern of viral entry occurring via endocytosis in cardiac cells.

We next explored the upstream regulators of the SARS-CoV-2 signaling response to identify molecular targets important for SARS-CoV-2 replication. By mapping reported kinase-substrate relationships to our phosphoproteomics data, we found that 37 kinases had substrates that were enriched for regulated phosphosites (Figure S6B, Figure 4I). We hypothesized that kinases with substrates enriched in up-regulated phosphorylation sites had increased activity in the SARS-CoV-2 response, and *vice versa* for kinases with down-regulated sites. The unique signaling responses of cardiac and lung cells was also evident at the kinase level, as of the 37 kinases enriched in at least one condition, 6 were significantly enriched in the same direction in both cell types. For kinases such as Protein Kinase C (PKC), some isoforms had opposite cell type-specific patterns of regulation. Of the PKC isoforms, PKCD had substrates enriched in upregulated sites in the lung cells, while PKCT was enriched in the cardiac cells (Figure 4I). This could suggest some convergence in signaling outcomes despite differing proteins being employed. However, the gold standard PKC substrate MARCKS S170 was uniquely upregulated in cardiac cells (Figure 4J).

Since SARS-CoV-2 proteins can also be phosphorylated by host kinases, we also searched our phosphoproteomics data specifically for phosphorylation of SARS-CoV-2 viral proteins. We measured 32 sites on 5 viral proteins (ORF1a, S, M, N and ORF9b (Table S1). This list includes S206 on the nucleocapsid protein, which has been reported as an SRSF Protein Kinase 1 (SRPK1) substrate (Figure 4K, (33)).

### Differential antiviral activity of candidate compounds in cardiac and lung AT2 cells

Based on the different transcriptional and phosphoproteomics data observed in lung and cardiac cells, we hypothesized that selected compounds would inhibit SARS-CoV-2 replication differentially. We interrogated our molecular datasets to predict new and alternative druggable targets. We selected 9 pathways that were differentially phosphorylated in SARS-CoV-2 infected lung AT2 and cardiac cells and identified 15 compounds that could inhibit the kinases critical for these pathways. These kinases (and their inhibitors) included: LIM domain kinase (R-10015), Checkpoint Kinases 2 (CCT241533 HCL), SRSF Protein Kinase 1 (Alectinib, SPHINX31), Cyclin-dependent Kinases (CDKs) (CCT251545, SNS-032, Palbociclib hydrochloride, Flavopiridol, Dinaciclib, Abemaciclib, Samuraciclib hydrochloride hydrate, Trilaciclib hydrochloride), Glycogen synthase kinase-3 (AZD1080) and protein kinase C (Bisindolylmaleimide I). We also tested an inhibitor of the TGFβ pathway using an inhibitor of the TGFβ type II receptor (GW788388). We screened the compounds at concentrations of 1, 10 and 50 μM for antiviral activity (data not shown) based on which we eliminated three compounds that targeted CDKs based on their strong toxicity in cardiac cells and lack of activity in lung cells (Flavopiridol, Dinaciclib, Samuraciclib hydrochloride hydrate). SNS-032 was analyzed further although it was toxic in cardiac cells because it showed promising antiviral activity in lung AT2 cells. We evaluated the remaining 12 compounds across a dose range from 0.04 to 30 μM for antiviral activity and cell cytotoxicity in both cardiac (H9 and NKX2-5)) and lung AT2 (H9) cells. Antiviral activity (% inhibition compared to the Vehicle control) was tested against nanoluciferase (nLuc)-expressing SARS-CoV-2 and an ancestral strain of SARS-CoV-2 (VIC01) by measuring luciferase expression and E gene copies, respectively. NHC was used as a positive control in all the antiviral assays

While the CDK8 inhibitor CCT251545 was ineffective in limiting SARS-CoV-2 replication (Figure S7), CDK4/6 inhibitors Abemacicilib, Palbociclib and Trilacicilib restricted virus replication in at least one cell type (Figure 5A). Abemaciclib and Palbociclib inhibited virus replication in both cell types, while Trilaciclib hydrochloride inhibited in cardiac cells with variable inhibition in the lung AT2 cells (Figure 5A). CDK2/7/9 inhibitor SNS-032 showed significant antiviral activity with minimal cytotoxicity in lung AT2 cells but was cytotoxic in cardiac cells (Figure 5A). In the lung AT2 cells, CHK inhibitor CCT241533HCl and PKC inhibitor Bisindolylmaleimide 1 showed some antiviral activity at the highest concentrations tested but were cytotoxic at these concentrations (Figure 5B, C). In cardiac cells, antiviral activity was observed with the CHK and PKC inhibitors at concentrations that were not cytotoxic (Figure 5B, C).

**Figure 5.**
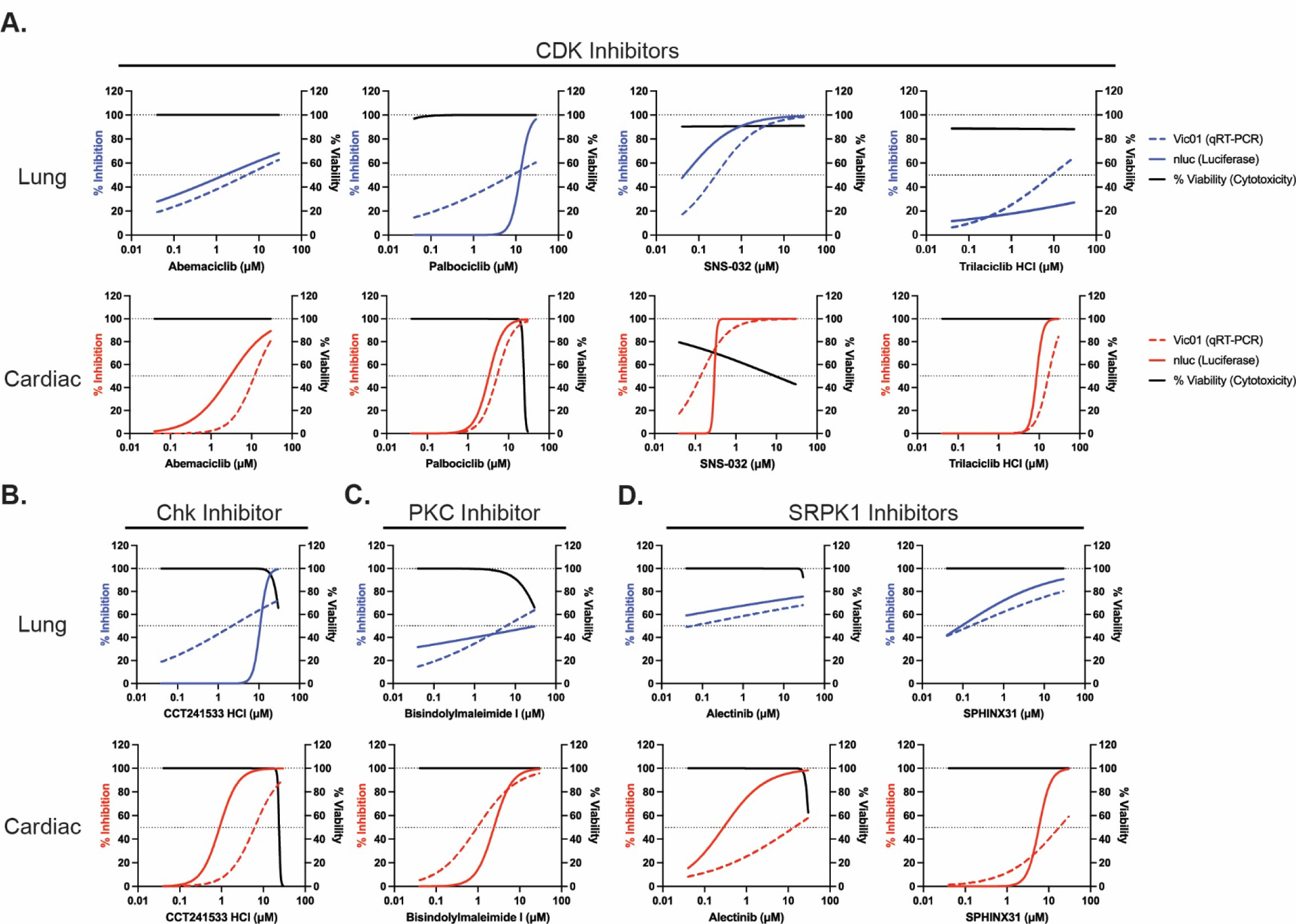
Efficacy of kinase inhibitors against SARS-CoV-2 replication varies between Lung AT2 and cardiac cells. Cell viability (% relative to Vehicle control, black lines) and inhibition of SARS-CoV-2 growth (% relative to Vehicle control) in the presence of CDK (A), CHK (B), PKC (C) and, SRPK1 (D) inhibitors in lung AT2 cells (H9) and cardiac cells (NKX2-5) at 2 dpi. Dotted black lines indicate 50% virus inhibition or cell viability.

Alectinib and SPHINX31, which inhibit SRPK1, inhibited viral replication in both lung AT2 and cardiac cells (Figure 5D). Our data suggest that while SRPK1 inhibitors can restrict viral replication in both cell types, greater inhibition was observed in lung AT2 cells. Similar activity of these antiviral drugs was observed in H9-derived cardiac cells (Figure S7A). Drugs that targeted LIM domain kinase (R-10015) and TGFβ type II receptor (GW788388) showed no activity in either cell type (Figure S7B). The GSK-3 inhibitor, AZD1080, showed no antiviral activity in H9-derived lung AT2 and cardiac cells (Figure S7). However, it did show antiviral activity in NKX2-5 cardiac cells against VIC01 but not the nLuc virus suggesting that removal of ORF7a may affect drug sensitivity (Figure S7,(2)). Overall, these data show that inhibition of kinases with small molecular inhibitors can abrogate SARS-CoV-2 replication with differing effectiveness in cardiac and lung cells.

## Discussion

Our data from parallel evaluation of SARS-CoV-2 infection in stem cell-derived cardiac and lung AT2 cells provide valuable insights into virus-host interactions in tissues that are significantly affected in COVID-19, with implications for rational design of therapeutic interventions. The key take-home messages from our study are that virus entry, cellular response, antiviral activity and cytotoxicity differ in SARS-CoV-2 infected human cardiac and lung AT2 cells and both differ from what is seen in African monkey kidney derived Vero cells that are widely used for SARS-CoV-2 research. Virus entry in both human cell types is dependent on ACE2 but further processing of the S protein is mediated through TMPRSS2 in lung AT2 cells, while infection of cardiac cells is achieved through the endosomal pathway and Cathepsin L. Host responses are significantly different, with a strong interferon response following infection in cardiac cells but not in lung AT2 cells. Phosphoproteomic analysis identified activation of different pathways in cardiac and lung AT2 cells. Parallel evaluation of antiviral activity and cytotoxicity of drugs in cardiac and lung AT2 cells reveals several points to consider in COVID-19 therapy, including the use of drug combinations to target both membrane protease and endosomal entry pathways, drugs that target SRPK1 and CDKs, and the importance of using relevant human stem-cell derived models in place of immortalized cell lines such as Vero cells for assessment of antiviral drugs.

Early reports characterizing SARS-CoV-2 demonstrated the requirement of ACE2 for virus entry (34). Our RNA-seq and qPCR analysis of lung and cardiac cells demonstrated high ACE2 expression in lung AT2 cells and low expression in cardiac cells. Immunofluorescence analysis of ACE2 expression in lung AT2 cells showed that only a small proportion of cells expressed ACE2, consistent with previous reports showing heterogeneous ACE2 expression in iPSC-derived and adult AT2 cells (24, 35). Despite low ACE2 expression in cardiac cells, virus replication was robust in this system achieving titres of more than 100,000 TCID_50_/mL by days four and six post-infection. Furthermore, SARS-CoV-2 infection of ACE2 KO lung AT2 and cardiac cells completely abolished virus growth, demonstrating that ACE2 is required for infection of these cells, consistent with previous reports using similar models (17, 36–38). Moreover, we showed that SARS-CoV- 2 infection was blocked with a combination of anti-ACE2 antibodies. Anti-ACE2 antibodies at a low dose did not completely block infection in AT2 cells but did in cardiac cultures, likely reflecting the difference in ACE2 expression between the cell types. Overall, we show that ACE2 expression is required for SARS-CoV-2 infection in stem-cell derived lung AT2 and cardiac cells.

To investigate the protease requirements for entry into stem-cell derived AT2 and cardiac cells, we performed entry inhibition assays with the TMPRSS2 inhibitor Camostat mesylate and the Cathepsin B/L inhibitor CA-074 Me (39). Camostat inhibited SARS-CoV-2 infection in AT2 cells, suggesting that TMPRSS2 is required for SARS-CoV-2 infection of AT2 cells, consistent with previous reports (24, 40). In contrast, CA-074 Me blocked SARS-CoV-2 infection in cardiac cells, demonstrating that Cathepsin L cleavage through endosomal entry is required for infection of cardiac cells, consistent with previous studies (17, 37, 38). The ability of SARS-CoV-2 to enter cells through different pathways may explain why clinical trials of Camostat mesylate in COVID-19 hospitalized patients and hydroxychloroquine alone did not result in clinical benefit (41). Our data suggest that a combination of drugs that target both pathways may be more effective *in vivo* and emphasize the importance of antiviral testing in several relevant tissue types. Ou *et al* reported that 10uM Camostat in combination with 10uM Hydroxychloroquine reduced virus replication compared to Camostat alone in Calu3 cells (42) and a clinical trial of Hydroxychloroquine in combination with Camostat is registered at clinicaltrials.gov (NCT04355052).

We found a robust interferon response in cardiac cells, which was not observed in lung AT2 cells. Induction of interferon in infected cardiac cell cultures has been previously reported using bulk RNA-seq (36, 38). However, Perez-Bermejo *et al* reported that single cell RNA-seq of infected iPSC-derived cardiomyocytes showed an induction of proinflammatory cytokines, but not type I or III interferons (17). A potential explanation for the discrepancy between our findings and those of Perez-Bermejo *et al* is that the additional cells present in our cultures (fibroblasts, smooth muscle cells and endothelial cells, (43)) or bystander cardiomyocytes are responsible for inducing robust interferon responses.

In contrast to cardiac cells, SARS-CoV-2 infection did not induce an interferon response in lung AT2 cells, consistent with another study of stem-cell derived AT2 models (44). The difference in host response to infection between cardiac cells and lung AT2 cells may be explained by the viral entry pathways. SARS-CoV-2 replication leads to detection of double stranded RNA (dsRNA) replication intermediates by Melanoma-differentiation-associated gene 5 (MDA5) and induction of a robust interferon response through Mitochondrial Antiviral Signaling Protein (MAVS) signaling (44, 45). Several studies have identified SARS-CoV-2 proteins (nsp1, nsp5, nsp6, nsp13, nsp15, ORF6 and ORF7b) that inhibit MAVS-induced Type I and III interferon responses (46–48). This mechanism of interferon antagonism likely explains the lack of interferon induction seen in lung AT2 cells following SARS-CoV-2 infection. In contrast, endosomal entry of SARS-CoV-2 in cardiac cells may explain the robust induction of interferon. Endosomes contain toll-like receptors (TLRs) including TLR3 and TLR7/8 that sense dsRNA and ssRNA, with only TLR3 detectable in cardiomyocytes (49). Furthermore, defects in TLR3 have been associated with disease severity in patients (50), suggesting TLR3 may play an important role in inducing interferon responses.

Although COVID-19 vaccines are highly effective in preventing severe illness and death, antiviral compounds are required for the treatment of COVID-19, particularly with the emergence of variant viruses and reduced effectiveness of vaccines in preventing symptomatic illness caused by variants of concern. To date, only a handful of drugs including Remdesivir, Molnupiravir and Paxlovid have been approved for use in hospitalized COVID-19 patients and additional drugs are needed. In this study, we established two cellular models relevant to COVID-19 disease that are scalable and amenable to high throughput screening. We used this system to screen drugs that advanced to clinical trials based on *in vitro* activity against SARS-CoV-2. We noted poor antiviral activity of Remdesivir in Vero cells compared to stem cell-derived lung AT2 cells and cardiac cells. These data are consistent with a previous report showing Remdesivir is metabolized inefficiently in Vero cells (51) and suggests that Vero cells are not optimal for screening antiviral compounds against SARS-CoV-2.

Based on results from phosphoproteomics analysis, we screened several kinase inhibitors for their ability to inhibit viral replication in lung AT2 and cardiac cells. Out of 12 compounds we evaluated, three had activity in both AT2 and cardiac cells (SPHINX31, Alectinib, Abemaciclib), one had activity in AT2 cells only (SNS-032), four had activity in cardiac cells only (CCT241533, Palbociclib, Trilaciclib, Bisindolylmaleimide I) and the remaining were ineffective. We observed that several CDK inhibitors were toxic at the concentrations tested, particularly in cardiac cells. Of note SNS-032 showed strong antiviral activity in lung AT2 cells but induced cardiac cell death; medicinal chemistry approaches could be explored to limit the toxicity for cardiac cells. Effectiveness of SRPK1 and CDK inhibitors against SARS-CoV-2 has been reported previously in Vero, Calu-3, A549-ACE2 cells and primary lung cells (33, 52). However, this is the first study to show potential toxicity of CDK inhibitors in cardiac cells. Although we observed antiviral activity in both cell types, SRPK1 inhibitors were effective at lower concentrations in lung AT2 compared to cardiac cells. Overall, we have identified several antiviral compounds with distinct effectiveness and toxicity profiles in lung AT2 and cardiac cells, highlighting the importance of using several cell types for evaluation of antiviral effectiveness.

In summary, with paired experiments in human stem cell derived tissue surrogates of organs that are affected in severe COVID-19, we have demonstrated important commonalities and several key differences in virus-host interactions in lung and cardiac cells. While the value of using human lung organoids, A549 and Calu-3 cells for evaluation of antiviral drugs has been recognized, our work highlights the importance of evaluation in additional relevant cells (lung and heart) for the evaluation of antiviral activity and cytotoxicity of drugs that are considered for treatment of COVID-19. This parallel evaluation in two cell types offers novel insights into rational design of therapeutic interventions.

## Acknowledgements

This work was supported by The Medical Research Future Fund (MRF9200007; K.S., E.R.P., J.M.P., D.A.E.) and the Victorian State Government (Victorian State Government (DJPR/COVID-19; KS, MR, ERP, J.P., D.A.E.) and COVID-19 Victorian Consortium (KS). E.R.P., M.R., D.A.E. are supported by The Novo Nordisk Foundation Center for Stem Cell Medicine (Grant NNF21CC0073729), The Stafford Fox Medical Research Foundation and The Royal Children’s Hospital Foundation. Fellowship support provided by National Health and Medical Research Council of Australia (E.R.P. and KS). MCRI is supported by the Victorian Government’s Operational Infrastructure Support Program. The Melbourne WHO Collaborating Centre for Reference and Research on Influenza is supported by the Australian Government Department of Health.

## Author Contributions

Conceptualization and design: E.P, J.P, S.J.H, M.R, D.E and K.S. Data acquisition: R.R, M.J.G, J.A.N, E.S.S, J.S, E.J.D, M.S, C.M-K, L.Y.Y.L, M.W, H.P, K.K and D.A-B. Data analysis and Interpretation: R.R, M.J.G, J.A.N, E.S.S, J.S, E.D, M.S, A.S, N.C, H.T.N and P.D.C. Reagent production: D.D and W-H.T. Manuscript preparation: R.R, M.J.G, J.A.N and K.S. Manuscript edits and proofing: E.S.S, J.S, E.J.D, M.S, E.P, J.P, S.H, M.R and D.E. All authors critically reviewed and approved the final version of the manuscript.

## Materials and Methods

### Cells

African green monkey kidney epithelial (Vero cells, ATCC Cat. CCL-81), Vero hSLAM (Merck, Car. 04091501), Calu-3 (ATCC, Cat. HTB-55) and VeroE6-TMPRSS2 (CellBank Australia, Cat. JCRB1819) cells were cultured at 37°C and 5% CO_2._ Vero cell media: Minimum Essential Media (MEM) (Media Preparation Unit, Peter Doherty Institute) supplemented with 5% Fetal Bovine Serum (FBS, Bovogen, Cat. SFBS), 50U/mL Penicillin and 50µg/mL Streptomycin (PenStrep, Thermo Fisher Scientific, Cat. 15070- 063), 2mM GlutaMAX (Thermo Fisher Scientific, Cat. 35050061) and 15 mM HEPES (Thermo Fisher Scientific, Cat. 15630130). Vero hSLAM cell media: MEM supplemented with 7% FBS, PenStrep, 2mM GlutaMAX, 15 mM HEPES and 0.4 mg/mL G418 Sulfate (Gibco, Cat. 10131027). Calu-3 cell media: MEM containing L-glutamine and sodium bicarbonate (Sigma, Cat. M4655) supplemented with 10% FBS, PenStrep, 1x non-essential amino acids (Gibco, Cat. 11140050) and sodium pyruvate (Fisher Scientific, Cat. BP356-100). VeroE6-TMPRSS2 cell media: Dulbecco’s Minimum Essential Media (DMEM) (Media Preparation Unit, Peter Doherty Institute) supplemented with 10% FBS, PenStrep, 2mM GlutaMAX and 1 mg/mL G418 Sulfate.

### Viruses

SARS-CoV-2 viruses hCoV-19/Australia/VIC01/2020 (VIC01, GISAID ID: EPI_ISL_406844), hCoV-19/Australia/VIC17991/2020 (Alpha variant, GISAID ID: EPI_ISL_779606), hCoV-19/Australia/QLD1520/2020 (Beta variant, GISAID ID: EPI_ISL_968081), hCoV-19/Australia/VIC18440/2021 (Delta variant, GISAID ID: EPI_ISL_1913206), hCoV-19/Australia/NSW-RPAH-1933/2021 (Omicron BA.1 variant, GISAID ID: EPI_ISL_6814922) and hCoV-19/Australia/VIC35864/2022 (Omicron BA.2 variant, GISAID ID: EPI_ISL_8955536) were a kind gift obtained from the Victorian Infectious Diseases Reference Laboratory (VIDRL). hCoV-19/Japan/TY7-503/2021 (Gamma variant, GISAID ID: EPI_ISL_877769, NR-54982) was obtained through BEI Resources, NIAID, NIH, contributed by National Institute of Infectious Diseases. icSARS-CoV-2-nLuc virus was a kind gift from Prof Ralph S. Baric from the Department of Microbiology and Immunology, University of North Carolina at Chapel Hill, Chapel Hill, NC, USA (2). SARS-CoV-2 VIC01 was propagated in Vero and Vero hSLAM cells in Vero infection medium (serum-free MEM in the presence of 1µg/mL TPCK-Trypsin (Cat. LS003740)). SARS-CoV-2 Alpha, Beta, Gamma and Delta variants were propagated in Vero hSLAM cells in infection medium (serum-free MEM with 1µg/mL TPCK-Trypsin). SARS-CoV-2 Omicron BA.1 and BA.2 variants were passaged in Calu3 cells in infection media (MEM containing 2% FBS). Virus stocks were stored at -80°C and titered as described below.

### Virus Titration

Virus titrations were performed in 96 well plates with confluent Vero and VeroE6-TMPRSS2 monolayers. Cells were washed with plain MEM and replaced with 180 µL of serum-free media containing 1µg/mL TPCK-Trypsin. Each sample was titrated in quadruplicate by adding 20µL of supernatant to first well and performing 10-fold serial dilutions. Cells were incubated at 37°C _and_ assessed microscopically for SARS-CoV-2- induced cytopathic effect (CPE) on day 4. Virus titres are expressed as mean log_10_TCID_50_/mL.

### Viral RNA extraction and RT-PCR

RNA was extracted as per the manufacturer’s recommendation using QiaCube HT (Qiagen) and QiaAmp 96 Virus QiaCube HT kit (Qiagen, Cat. 57731). RT-PCR reaction was setup using SensiFast Probe No-ROX One- Step Kit (Bioline, Cat. BIO-76005) using the following primers/probes: E_Sarbeco_F1: 5’- ACAGGTACGTTAATAGTTAATAGCGT-3’, E_Sarbeco_R2: 5’-ATATTGCAGCAGTACGCACACA-3’), E_Sarbeco_P1_FAM: 5’-ACACTAGCCATCCTTACTGCGCTTCG-3’. Serial 10-fold dilutions of plasmid encoding the viral E gene were used to generate a standard curve for calculating the virus genome copies in the samples.

### Lung alveolar type 2 cell differentiation

Induced pluripotent H9 (female) stem cells were seeded onto flasks coated with Matrigel (Corning, Cat. 354230) in Essential 8 medium (Thermo Fisher Scientific, Cat. A1517001). After 48 h, medium was changed daily with RPMI 1640 (Thermo Fisher Scientific, Cat. 21870084) supplemented with B-27 (Gibco, Cat. 17504044), 100ng/mL Activin A (Peprotech, Cat. 120-14P), 1µM CHIR99021 (Sigma-Aldrich, Cat. SML1046), PenStrep for 3 days. On days 4-8, medium was changed daily with DMEM/F12 media (Thermo Fisher Scientific, Cat. 105650) supplemented with N2 (Gibco, Cat. 17502048), B27, 0.05mg/mL Ascorbic Acid (Sigma-Aldrich, Cat. A92902), 0.4mM Monothioglycerol (Sigma-Aldrich, Cat. M6145), 2µM Dorsomorphin (Stemcell Technologies, Cat.72102), SB431542 (Miltenyi Biotec, Cat. 130-106-543) and PenStrep. On days 9-12, medium was changed daily with DMEM/F12-based medium with B27, 0.05mg/mL Ascorbic Acid, 0.4mM Monothioglycerol, 20ng/mL BMP4 (Peprotech, Cat. 120-05ET), 0.5µM Retinoic Acid (ATRA) (Sigma-Aldrich, Cat. R2625), 3µM CHIR99021, PenStrep. Day 12 and onwards, medium was changed every other day with DMEM/F12 supplemented with B27, 0.05mg/ml Ascorbic Acid, 0.4mM Monothioglycerol, 10ng/mL FGF10 (Stemcell Technologies, Cat. 78037), 10ng/mL FGF7 (Peprotech, Cat. 10019), 3µM CHIR99021, 50nM Dexamethasone (Sigma-Aldrich, Cat. D4902), 0.1mM 8- Bromoadenosine 3′,5′-cyclic monophosphate (8-Br-cAMP) (Sigma-Aldrich, Cat. B5386), 0.1mM 3-Isobutyl-1-methylxanthine (IBMX) (Sigma-Aldrich, Cat. I5879) and PenStrep. Cultures were embedded onto Matrigel on day 18 in 12-well plates. On day 30, organoids were dissociated in TrypLE (Thermo Fisher Scientific, Cat. 12604013) for 3 mins before re-embedding in Matrigel. Lung organoids were maintained for experiments between passage 2-8 prior to dissociation with TrypLE and seeding onto Geltrex (Gibco, Cat. A1413201) -coated plates supplemented with Y-27632 (Selleck Chemicals, Cat, S1049) at the initial seeding step. Cells were maintained for a further 7-10 days in 2D culture until infection at 70%+ confluency.

### Cardiac Cell Differentiation

The human embryonic stem cell lines HES3 *NKX2-5^eGFP/w^* and H9 (both female), and human induced pluripotent stem cell line MCRIi010-A (male), were used for viral infection studies in 2D monolayer cultures. Each stem cell line and their derivatives were cultured as outlined previously (30). Cardiomyocytes cultures were differentiated as previously described and cryopreserved at day 10 following differentiation (27). Cells were subsequently thawed in basal differentiation media containing RPMI 1640 (Thermo Fisher Scientific, Cat. 21870), 2% B27 without vitamin A (Thermo Fisher Scientific, Cat. 12587), 1% Glutamax (Thermo Fisher Scientific, Cat. 35050), 0.5% PenStrep (Thermo Fisher Scientific, Cat. 15070) and 10 µM Y-27632 for 24 hours at 37⁰C. Cells were then maintained in basal differentiation medium for an additional 2 days. To enrich for cardiomyocytes, cells were cultured for 2 days in lactate purification media as described previously (53). The cells were maintained in maturation media, as described previously (43), from day 15 to day 23 post differentiation prior to viral infection.

### CRISPR/Cas9 ACE2 KO line generation

For CRISPR/Cas9 knock-out of *ACE2*, synthetic oligonucleotides containing sgRNAs (5’ sgRNA 5’- CACCGTCTAGGGAAAGTCATTCAG-3’ and 5’- AAACCTGAATGACTTTCCCTAGAC-3’ and 3’ sgRNA 2 5’- CACCGCAGTAATCTAATCTTTAAG-3’ and 5’-AAACCTTAAAGATTAGATTACTGC-3’ targeting the first coding exon of *ACE2* were generated with overhangs for BbsI digestion and subsequent cloning into pX458 (Addgene, Cat. 48138) as described previously. Cells were harvested with TrypLE, and transfections were performed using the Neon Transfection System (Thermo Fisher Scientific). Electroporation was performed in a 100 μL tip using the following conditions: 1050 V, 30 ms, 2 pulses. Following electroporation, cells were transferred to flasks containing mitotically inactivated MEFs and HES or iPSC media (without PenStrep) with 5 μM Y-27632, which was omitted in subsequent media changes. Cells transiently expressing EGFP were single cell sorted, colonies were grown for 2 weeks and screened via PCR.

### Clone characterization: karyotyping, pluripotency, and differentiation

Genomic integrity of ACE2 KO lines was assessed using the Illumina Infinium GSA-24 v2.0 and was performed by the Victorian Clinical Genetics Service, Royal Children’s Hospital (Melbourne). Pluripotency was examined via expression of EPCAM-BV421 (Biolegend, Cat. 324220), CD9-FITC (BD Biosciences, Cat. 555371) and SSEA4-PE/Cy7 (Biolegend, Cat. 330420) expression using flow cytometry. IgG-BV421, IgG-FITC and IgG-PE/Cy7 (Biolegend, Cat. 400158 and BD Biosciences Cat. 555748, 400126) were used for isotype controls. Cardiac cultures generated from the WT and ACE2 KO lines were then assessed using cardiomyocyte specific markers, cardiac troponin T and α-Actinin (Abcam, Cat. AB45932, A7811) to demonstrate comparable levels of cardiomyocyte differentiation. AF488 and AF647 (Thermo Fisher Scientific Cat. A11034, A21236) were used for detection on the LSR Fortessa X-20 Flow cytometer (BD Biosciences).

### Quantitative PCR

Total RNA was extracted using Trizol reagent (Thermo Fisher Scientific, Cat. 15596026) following the manufacturer’s protocol or using the RNeasy Mini Kit (Qiagen, Cat. 74106). cDNA synthesis was conducted using the SuperScript™ VILO™ Master Mix (Thermo Fisher Scientific, Cat. 11755050) according to the manufacturer’s instructions. Quantitative real-time PCR (qPCR) was performed with either TaqMan Fast Advanced Master Mix, TaqMan™ Gene Expression Master Mix or PowerUp™ SYBR™ Green Master Mix (Thermo Fisher Scientific, Cat. 4444557, 4369510 or A25777 respectively). The Taqman probes (Thermo Fisher Scientific, Cat. 4453320) are listed in Table S3. Green are as follows: *ACE2* 5’-TTAACCACGAAGCCGAAGAC-3’ and 5’- TACATTTGGGCAAGTGTGGA-3’; 5’-CCTGGCTGAAAGACCAGAAC-3’ and 5’- GCAACAGATGATCGGAACAG-3’; and 5’- GGTTGGCATTGTCATCCTG-3’ and 5’ GGAGGTCTGAACATCATCAGTG-3’. Glyceraldehyde-3-phosphate dehydrogenase (*GAPDH*) (5’ TGCACCACCAACTGCTTAGC-3’ and 5’ GGCATGGACTGTGGTCATGAG-3’) was used as a housekeeping gene for normalization.

### Western blot analysis

hESC and iPSC-derived cardiac and lung AT2 cells were lysed in high salt RIPA buffer (RIPA+0.5M NaCl) with added Protease inhibitor (Sigma-Aldrich, Cat. 5056489001) and phosSTOP (Roche, Cat. 4906845001). Lung AT2 lysates were made using Laemmli sample buffer (Bio-Rad, Cat: #1610737EDU). Total protein concentration was measured by BCA assay (Thermo Fisher Scientific, Cat. 23227). 20 µg of samples were loaded into a 4–20% Mini-PROTEAN pre-cast gel (Bio-Rad, Cat: 4561093) and ran at 70 V for 30 min, followed by ∼70 min at 120 V. Protein was transferred to PVDF membrane and blocked in 5% skim milk in Tris-buffered saline contain Tween-20 (TBST) for 1 h. The membrane was incubated with primary antibodies against ACE2 (Abcam, Cat. 15348, 1:1000 in 5% skim milk overnight at 4⁰C) and GAPDH as a loading control (Fitzgerald Industries International, Cat. 10R-2932, 1:3000 in 5% skim milk/TBST for 1 h at room temperature (RT)). HRP-linked antibodies (Bio-Rad, Cat. 1706516 and Cell Signaling Technology, Cat. 7074S) were then used at 1:3000 in skim milk/TBST for 1 h at RT. Blots were imaged on a GE Amersham Imager 680 following addition of ECL substrate (Bio-rad, Cat. 170-5060).

### Virus growth in lung AT2, cardiac and Vero cells

Infection of human ESC and iPSC-derived cells and Vero cells was performed in 24 well tissue culture plates. Vero cells were washed with MEM prior to infection. Media was removed and replaced with 10^4^ TCID_50_ of SARS-CoV-2 in 100ul and incubated for 1 h at RT. The inoculum was removed, cells washed twice with cell-specific media and refed with media. The second wash was harvested for day 0 sampling. Vero cells were cultured in serum-free media containing 1µg/mL TPCK-Trypsin. Supernatants were collected each day and media was replenished. Harvested supernatants were stored at -80°C to determine infectious virus titres and E gene copies.

### Immunofluorescence

Cells were fixed with 4% PFA and blocked with blocking buffer (1% horse serum and 0.1% Triton X-100 in phosphate-buffered saline (PBS)) and stained with primary antibodies (cTNT: Thermo Fisher Scientific, Cat. MA1-16687, 1:200; dsRNA: Australian Bioresearch, Cat, ab01299-2.0, 1:200; SP-C: Santa Cruz Biotechnology, Cat. sc-518029,1:100; AQP5: Santa Cruz Biotechnology, Cat. sc-514022, 1:100; NKX2.1: Abcam, Cat. ab76013, 1:100) overnight at 4⁰C. Cells were washed with PBS and stained with secondary antibodies from Invitrogen (Mouse IgG2a AF594 Cat. A21135, Mouse IgG1 AF647 Cat. A21240, Rabbit IgG AF555 Cat. A21428, Mouse IgG2b AF488 Cat. A21141, Mouse IgG2a AF555 Cat. A21127, Rabbit IgG AF647 Cat. 21244) and Hoechst 33342 (Life Technologies) at 1:1000 in blocking buffer for 1-2 hours at RT. For ACE2 staining, cells were blocked with 10% normal goat serum (Thermo Fisher Scientific, Cat. PCN5000) in PBSTT (PBS with 0.1% Triton-X and 0.1% Tween). Primary ACE2 antibodies (WCSL141, (23)) and cTNT antibodies were left to incubate overnight at 4⁰C in 5% goat serum in PBSTT before being washed in PBS and probed with secondary antibodies (Jackson Immuno, Cat. 709-116-098 at 1:100 and Hoescht at 1:200).

### SARS-CoV-2 entry inhibition

Lung AT2 and cardiac cells seeded into 24 well plates were treated with 100µL of α-ACE2 antibodies (WCSL141 and WCSL148, (23)), human IgG isotype control, Camostat Mesylate (Sigma Aldrich, Cat. SML0057) or DMSO for 1 h at 37°C. Subsequently, 10^4^ TCID_50_ of SARS-CoV-2 was added to cells and incubated for 1 h at 37°C. Virus inoculum was removed, and cells were washed with plain MEM twice before replacement with the 500µL cell-specific culture media. For CA-074 Me inhibition, cells seeded into 24 well plates were treated with 200µL of CA-074 Me (Selleck Chemicals, Cat. S7420) for 2 h at 37°C. Subsequently, 10^4^ TCID_50_ of SARS-CoV-2 was added to cells and incubated for 1 h at 37°C. Supernatant samples were obtained daily, and the media was replaced with drug-containing media until day 3. Infectious virus titres and E gene copies were determined as detailed above.

### Antiviral testing

Compounds remdesivir (MedChemExpress, Cat. HY104077), NHC (β-D-N4-hydroxycytidine, MedChemExpress, Cat. HY125033), favipiravir (Toyama Chemicals Co. Ltd, Japan, T705), tizoxanide (Romark Laboratories, Tampa, FL) chloroquine (Sigma Aldrich, Cat. C6628) and piperaquine (Sigma Aldrich, Cat. C7874) were tested in lung AT2, cardiac and Vero cells in 24 well tissue culture plates. Vero cells were washed with MEM prior to addition of 100µL of cell type-specific media containing diluted compounds. The vehicle controls were prepared to contain the same amount of vehicle (DMSO or water) as the 10µM of compound. One hour after addition of diluted compound, 100µL of media containing 10^4^ TCID_50_ of SARS-CoV-2 (and 1µg/ml TPCK-Trypsin for Vero cells) was added and incubated for an additional hour at RT. The inoculum was removed and replaced with 500µL of cell-specific media containing diluted compounds. At 3 dpi, supernatants were harvested and stored at -80°C. Infectious virus titres and E gene copies were determined as detailed above.

Kinase inhibitor compounds (Table S2) were tested in lung AT2 and cardiac cells seeded in 96 well culture plates. Cells were incubated with 200μL of cell-specific media containing 30μM to 0.04μM of compound for 2 h at 37°֯C. DMSO was maintained consistently to a final concentration of 0.1%. After 2 h, 10^4^ TCID_50_ of icSARS-CoV-2-nLuc or VIC01 was added to each well (20uL total) and cells were incubated at 37֯°C for 2 days. Cell supernatant from VIC01-infected cells was collected for measurement of E-gene copies as detailed above. For nLuc virus detection, cells were lysed with Passive Lysis Buffer (Promega, Cat. E1941) and luciferase expression measured using Nano-Glo Luciferase Assay System (Promega, Cat. N1130) as per the manufacturer’s instructions. Luminescence was measured on FLUOstar Omega (BMG Labtech).

### Antiviral toxicity testing

All compounds were tested in lung AT2, cardiac and Vero cells in 96 well tissue culture plates. Vero cells were washed with 200µL of MEM. After removal of media, 100µL of cell type-specific media containing diluted compound was added. At day 2 (Kinase inhibitors) or day 3 (other compounds), cell viability was measuring using the CellTiter-Glo®2.0 cell viability kit (Promega, Cat. G9241) as per manufacturer’s recommendations. Luminescence was measured on FLUOstar Omega (BMG Labtech).

### RNA Sequencing Analysis

Human stem cell-derived cardiac and lung cells grown in 24 well tissue culture plates were infected with 10^4^ TCID_50_ of SARS-CoV-2. At 0-, 1- and 3- days post-infection, supernatant was removed, and the cell monolayer was lysed with 500µL of TRIzol reagent (Life Technologies Australia PTY LTD, Cat. 15596018). RNA was extracted following the manufacturer’s protocol. RNA Sequencing data were demultiplexed using a modified version of the Sabre demultiplexer to produce a single fastq file per sample. Fastq files were processed using the RNAsik pipeline [https://doi.org/10.21105/joss.00583]. Reads were aligned to EnsEMBL GRCh38 (54) using the STAR aligner (55) and duplicates were marked with Picard [“http://broadinstitute.github.io/picard/”]. Aligned reads were quantified to gene level counts using featureCounts (56). Next, differential gene expression analysis was performed in Degust [https://doi.org/10.5281/zenodo.3501067] using the EdgeR QL method (57) to produce sets of differentially expressed genes for Cardiac and Lung conditions, respectively. The sets of differentially expressed genes were processed for pathway enrichment using Metascape (58) with default parameters except that only the WikiPathways ontology was used. Figures were generated using R [4] and tidyverse [https://doi.org/10.21105/joss.01686] packages. RNA-seq data have been submitted to the NCBI GEO database (GSE212003).

### LEGENDplex

Cytokine/chemokine concentrations in supernatant from mock and SARS-CoV-2 infected lung AT2 and cardiac cells were analyzed using the LEGENDplex human anti-virus response panel (Biolegend, Cat. 740390) following the manufacturer’s instructions. Samples were run on a BD FACSCanto II and analyzed using LEGENDplex™ Data Analysis Software Suite (v8).

### Phosphoproteomics

Human cardiac and lung AT2 cells grown in 6 well tissue culture plates inoculated with mock or SARS-CoV-2 were harvested at 0-, 18- and 24- hours post-inoculation for phosphoproteomic analysis. Briefly, media was removed and replaced with 500µL of respective media containing 5×10^4^ TCID_50_ of SARS-CoV-2. Mock wells received 500µL of media alone. After infection for 1 h, inoculum was removed and replaced with 2000µL of respective media. At selected timepoints post-infection, wells were washed four times with 5ml of ice-cold TBS. After TBS removal, cells were lysed with 100µL of lysis buffer (6M Guanidinium chloride, 100mM Tris pH 8.5), inactivated at 95°C for 5 minutes and frozen at -80°C. Upon thawing, lysates were sonicated with a tip- probe sonicator (50% output power, 30 seconds), and an aliquot was diluted 1:5 in 8M Urea to determine protein concentration by BCA assay. Protein (250 µg) was diluted in SDC buffer, reduced and alkylated at 45°C for 5 min by the addition of 10 mM Tris (2- carboxyethyl) phosphine (TCEP)/40 mM 2-Chloroacetamide (CAA) pH 8, and digested by the addition of 1:100 Lys-C and Trypsin overnight at 37°C with agitation (1,500 rpm). After digestion phosphopeptides were enriched in parallel using the high-sensitivity EasyPhos workflow as previously described (32). Eluted phosphopeptides were dried in a SpeedVac concentrator (Eppendorf) and resuspended in MS loading buffer (0.3% TFA/2% acetonitrile) prior to LC-MS/MS measurement.

Phosphopeptides were loaded onto a 55 cm column fabricated in-house, from 75μM inner diameter fused silica packed with 1.9μM C18 ReproSil particles (Dr. Maisch GmBH), and column temperature was maintained at 60°C using a Sonation column oven. A Dionex U3000 RSLC Nano HPLC system (Thermo Fisher Scientific) was interfaced with a Q Exactive HF X benchtop Orbitrap mass spectrometer using a NanoSpray Flex ion source (Thermo Fisher Scientific). Peptides were separated with a binary buffer system of 0.1% (v/v) formic acid (buffer A) and 80% (v/v) acetonitrile / 0.1% (v/v) formic acid (buffer B) at a flow rate of 400nL/min, and separated with a gradient of 3-19% buffer B over 40 min, followed by 19-41% buffer B over 20 min, resulting in a gradient time of 60 min. Peptides were analyzed in Data Independent Acquisition (DIA) mode, with one full scan (350-1,400 m/z; R = 120,000 at 200 m/z) at a target of 3e6 ions, followed by 48 DIA MS2 scans using HCD (target 3e6 ions; max. IT 22 ms; isolation window 14.0 m/z; NCE 25%, window overlap 1m/z), detected in the Orbitrap mass analyzer (R = 15,000 at 200 m/z). RAW MS data was processed using Spectronaut v15.4.210913.50606, with searches performed using the directDIA method against the Human and SARS-CoV2 UniProt databases (January 2021 and October 2021 releases respectively). Default “BGS Phospho PTM Workflow” settings were used, with a linear model used for PTM Consolidation, Cross Run Normalization enabled, and a localization Probability Cutoff of 0.5. Data filtering mode was set to ‘Qvalue’.

Phosphoproteomes were log transformed and median normalized. Ratios of SARS-CoV- 2 to the median mock values were taken for each timepoint in each cell type. Regulated phosphopeptides were identified with a one-way ANOVA for each cell type separately. P- values were adjusted for multiple hypothesis testing with the q-value R package. Dunnett’s post hoc tests were performed to determine the timepoint at which the regulation occurred, with the 0-hour timepoint as the control condition. Enrichment of regulated phosphosites was determined by ranking phosphosites by their Log2FC and performing a modified weighted gene set enrichment analysis, implemented in the R package ksea. Cellular component annotation was obtained from GO and reported kinase-substrate relationships were obtained from PhosphoSitePlus.

### Integration of RNA-seq and phosphoproteomics

To systematically identify and prioritize the drivers of the molecular response to SARS-CoV-2 infection, based on the combination of transcriptomics and phospho-proteomics data, we run the ExIR model separately on heart and lung datasets using the R package influential (https://cran.r-project.org/package=influential). ExIR is a versatile one-stop model for the extraction and prioritization of candidates from high-throughput data. In particular, we used the following three input data for running the ExIR model: 1) entire transcriptomic dataset as the experimental data, 2) table of differentially expressed genes, and 3) the list of differentially phosphorylated proteins at 72-hour time-point as the desired list of features. In this way, the ExIR model is built based on the differentially phosphorylated proteins in the context of the transcriptomic data. Lastly, the top five drivers prioritized by ExIR were visualized separately for lung and heart dataset.

### Statistical analysis

All data were plotted and analyzed using GraphPad Prism 9. Log_10_ virus titres and E gene copies were analyzed using either a student’s T-test or two-way ANOVA with Tukey’s or Dunnett’s multiple comparisons test as appropriate. *, p<0.05; **, p<0.01; ***, p<0.001 and ****, p<0.0001. Dotted lines indicate the lower limit of detection of the assay unless indicated otherwise. Data are representative of at least two independent experiments showing mean (±SD) unless indicated overwise. Antiviral activity and cytotoxicity data for the kinase inhibitors (Fig 5 and S7) was analyzed by calculating percent inhibition/viability relative to vehicle control. Curve fitting was performed using non-linear regression (four parameters – variable slope) with data constrained between 0 and 100.

## Figure Legends

**Figure S1.**
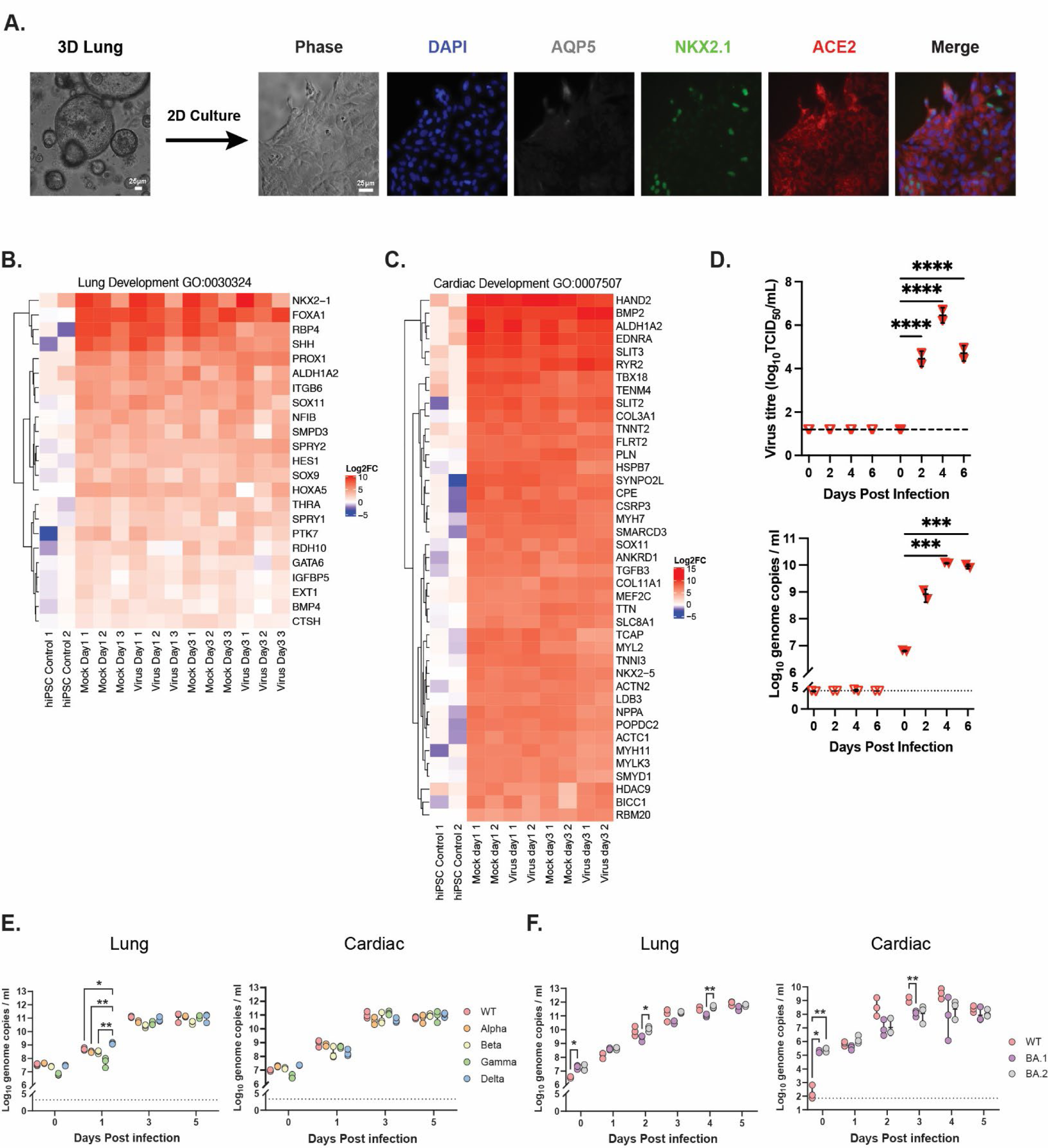
(A) Immunofluorescence images of lung culture from 3D organoids into 2D (AQP5, White; NKX2.1, Green; ACE2, Red). Heatmap analysis of lung-related (B) and cardiac-related (C) gene sets. (D) Viral titres and genome copies in supernatant from cardiac cells (ALK3-KO) infected with SARS-CoV-2 (VIC01). Viral genomes in supernatants from cardiac and lung AT2 cells infected with VIC01 (WT) compared to cells infected with Alpha, Beta, Gamma and Delta variants (E) or Omicron (BA.1 and BA.2) variants (F).

**Figure S2.**
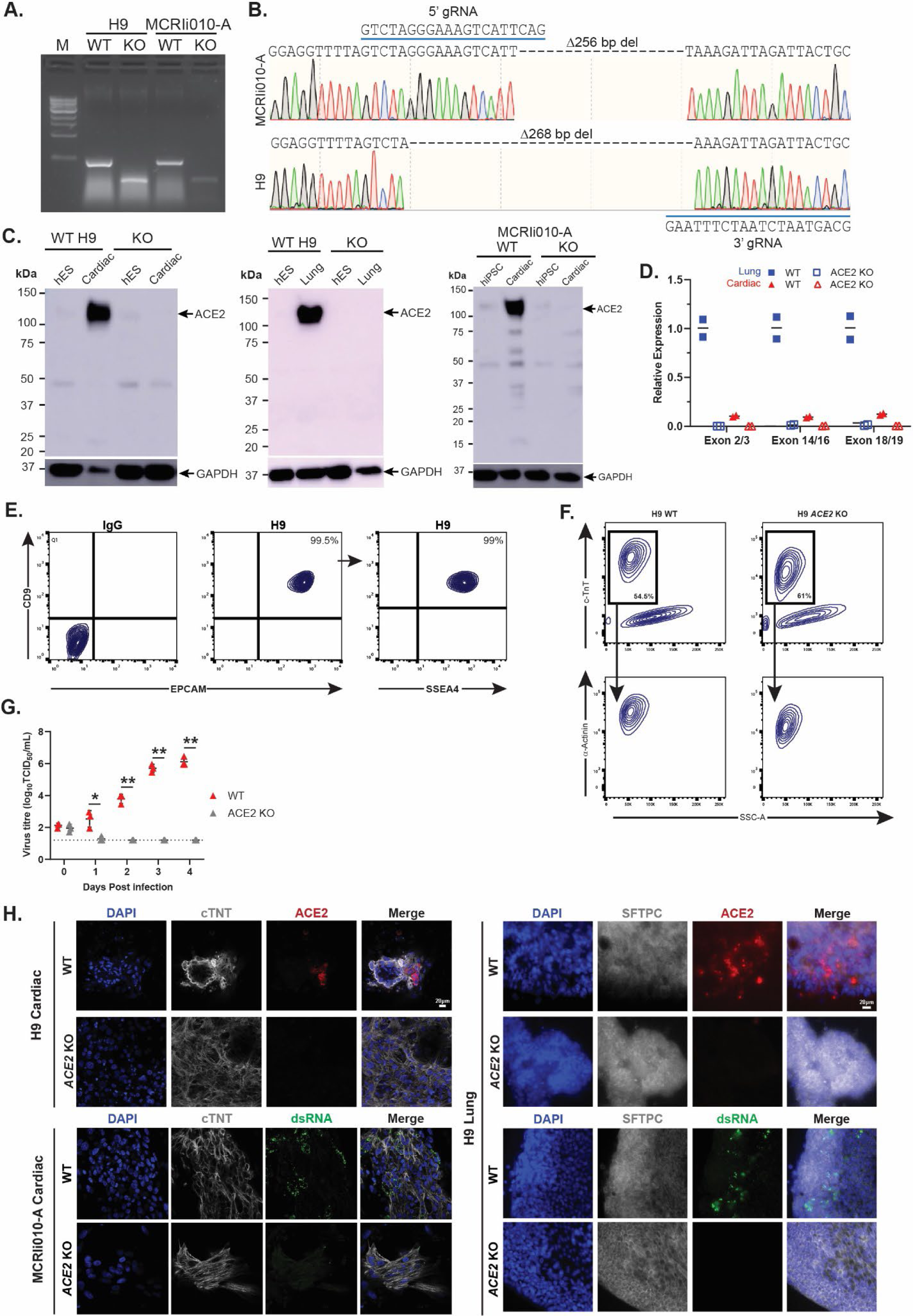
(A) PCR for *ACE2* in both the H9 and MCRIi010-A *ACE2* WT and KO clones. (B) Sanger sequencing of MCRIi010-A and H9 ACE2 KO clones. (C) ACE2 expression in ESCs/iPSCs and differentiated cardiac and lung AT2 cells. GAPDH was used as a loading control. (D) Relative expression of *ACE2* in the H9 ACE2 KO-derived cardiac and lung AT2 cells compared to lung AT2 WT. (E) Representative flow cytometry plot of proteins associated with pluripotency (CD9, EPCAM, SSEA4) in the H9 *ACE2* KO line. (F) Representative flow cytometry plot of cardiac proteins (c-TnT, α-actinin) following differentiation in H9 WT and ACE2 KO lines. (G) Virus titer in supernatant from SARS- CoV-2 infected ACE2 KO and WT MCRIi010-A-derived cardiac cells. (H) Representative fluorescent confocal microscopy images of ACE2 (red) and dsRNA (green) expression in WT and *ACE2 KO* cardiac (cTNT-positive) and lung AT2 (SFTPC-positive) cells (H9 and MCRIi010-A). Scale bars are highlighted on each panel.

**Figure S3.**
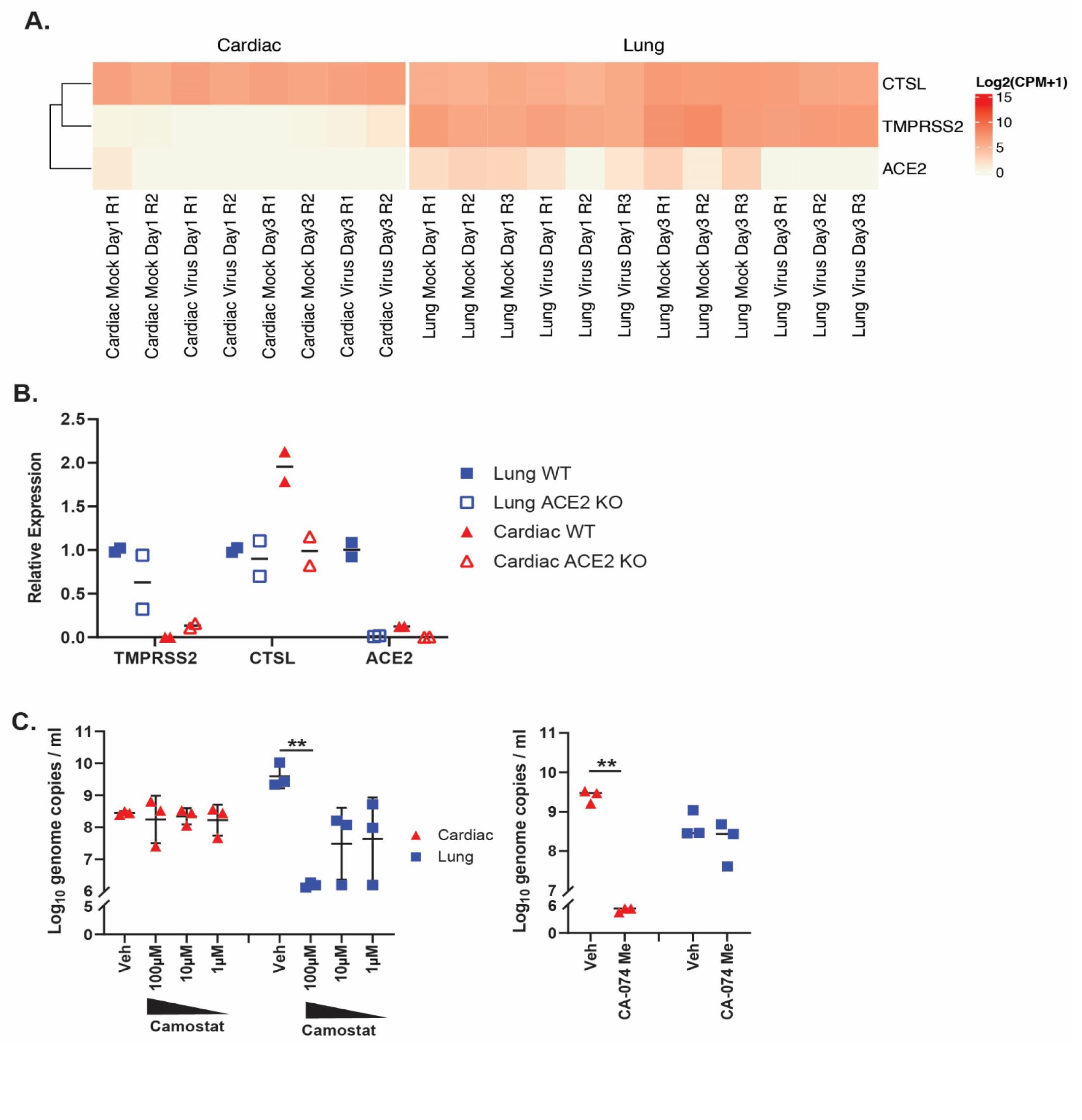
(A) Heat map in log2 counts per million (CPM) representing the gene expression of ACE2, TMPRSS2 and Cathepsin L (CTSL) in uninfected and SARS-CoV- 2-infected iPSC-derived lung AT2 and cardiac cells at 1 and 3 dpi. (B) Relative expression of ACE2, TMPRSS2 and Cathepsin L in iPSC-derived cardiac and lung AT2 ACE2 KO cells compared to lung AT2 WT. (C) Viral genome copies in supernatants from cardiac (red triangles) and lung AT2 (blue squares) cells at 3 dpi infected with SARS-CoV-2 in the presence of Camostat, CA-074 Me or DMSO.

**Figure S4.**
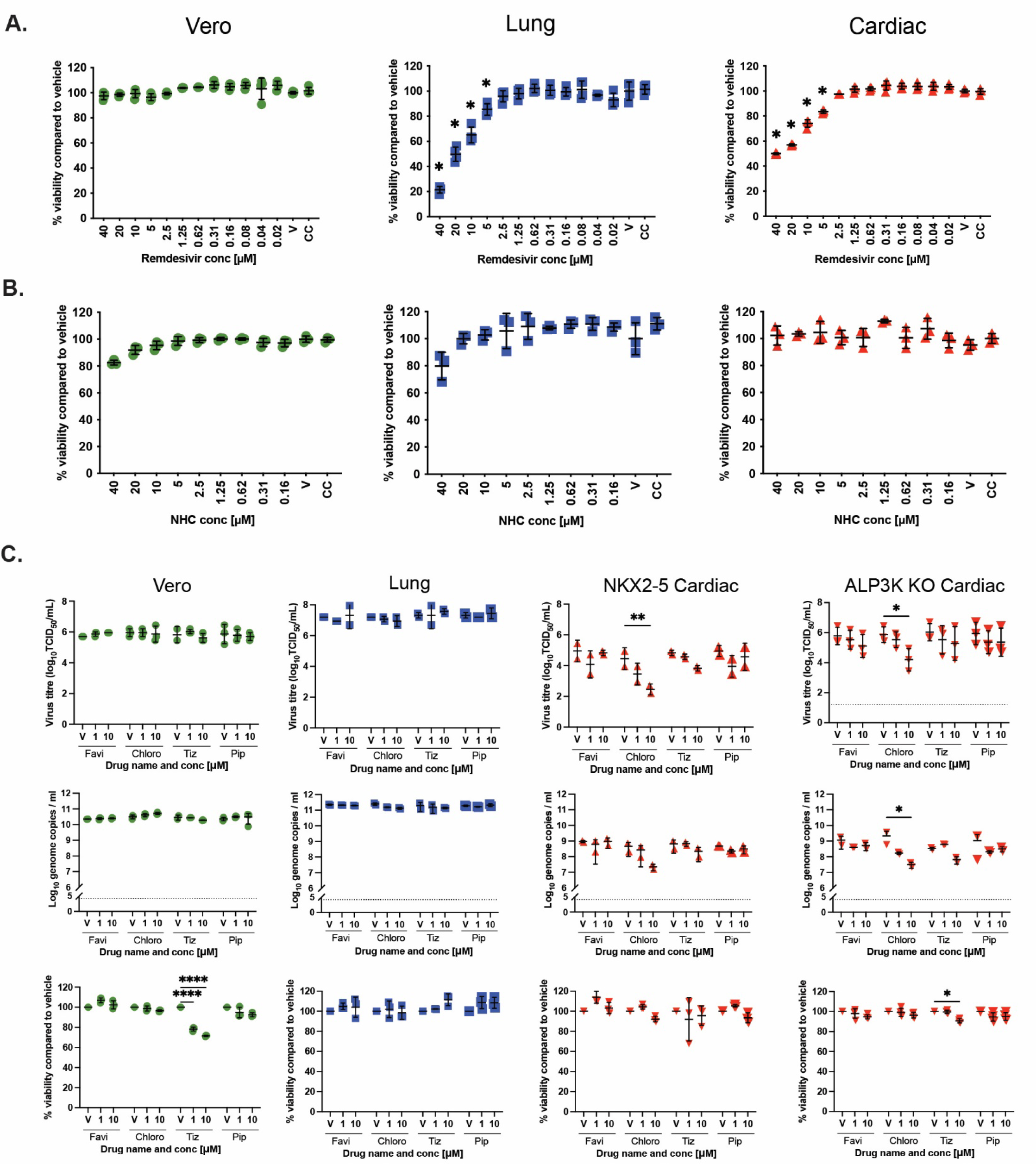
(A) Cell viability in uninfected Vero, lung AT2 and cardiac cells following 3 days culture in the presence of Remdesivir or NHC at the indicated concentrations. (B) Virus titer and viral genome copies in the supernatant at 3 dpi in Vero, lung AT2, NKX2- 5 cardiac and ALP3K KO cells infected with SARS-CoV-2 in the presence of 1µM or 10µM Favipiravir, Chloroquine, Tizoxanide or Piperaquine. Cell viability of uninfected cells after 3 days culture in the presence of the above drugs.

**Figure S5.**
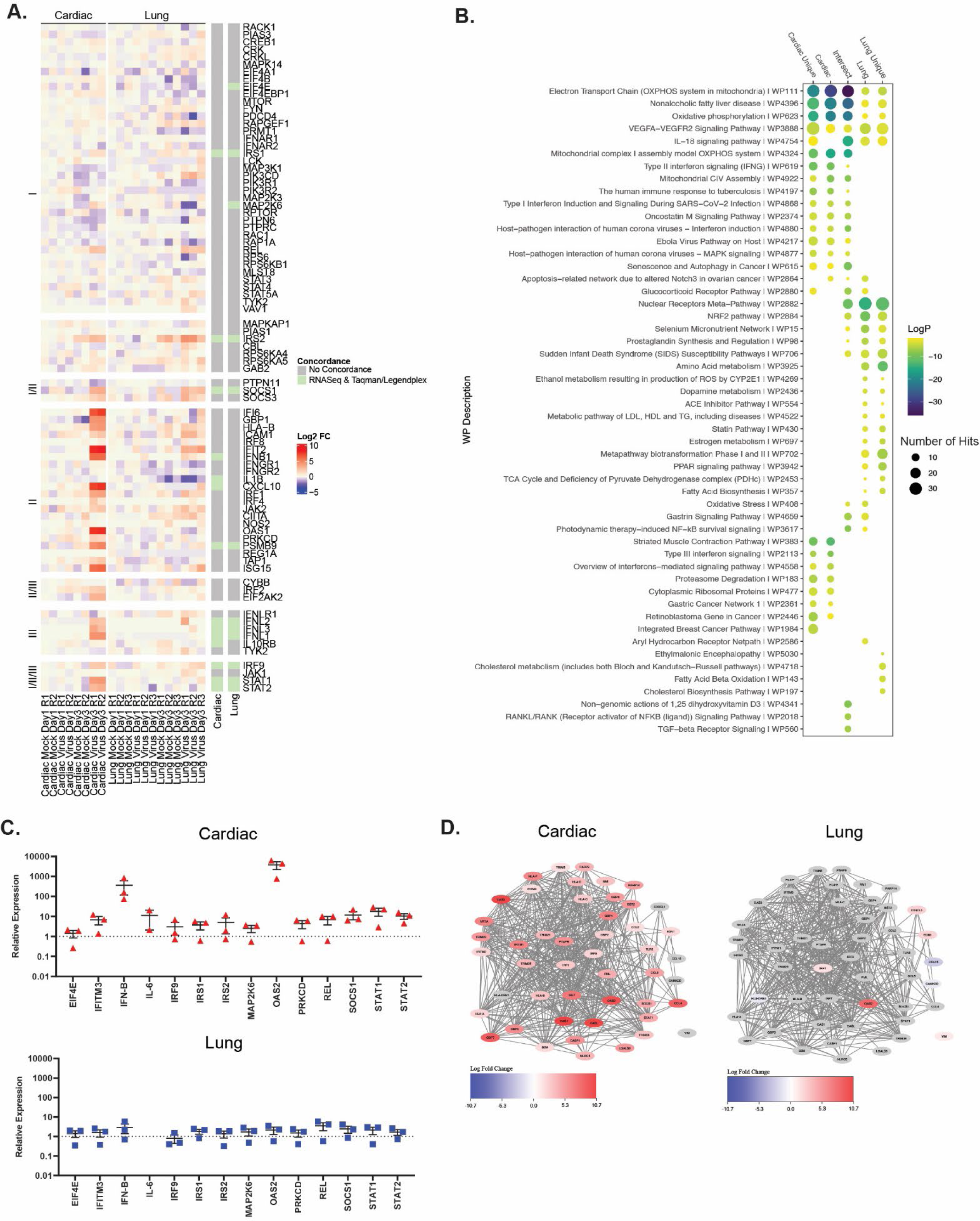
(A) Extended heat map in Log2 fold change (FC) of representative interferon genes (compared to representative mock-infected samples at 1 dpi) separated by type as defined by WikiPathways. (B) Extended Plot of top enriched WP terms. (C) Relative expression of IFN pathway genes compared to mock-infected cells. (D) Gene network diagram of the GO Term ’response to interferon gamma’ (GO:0034341). Node colors are log2 fold change of the gene in the respective tissue type.

**Figure S6.**
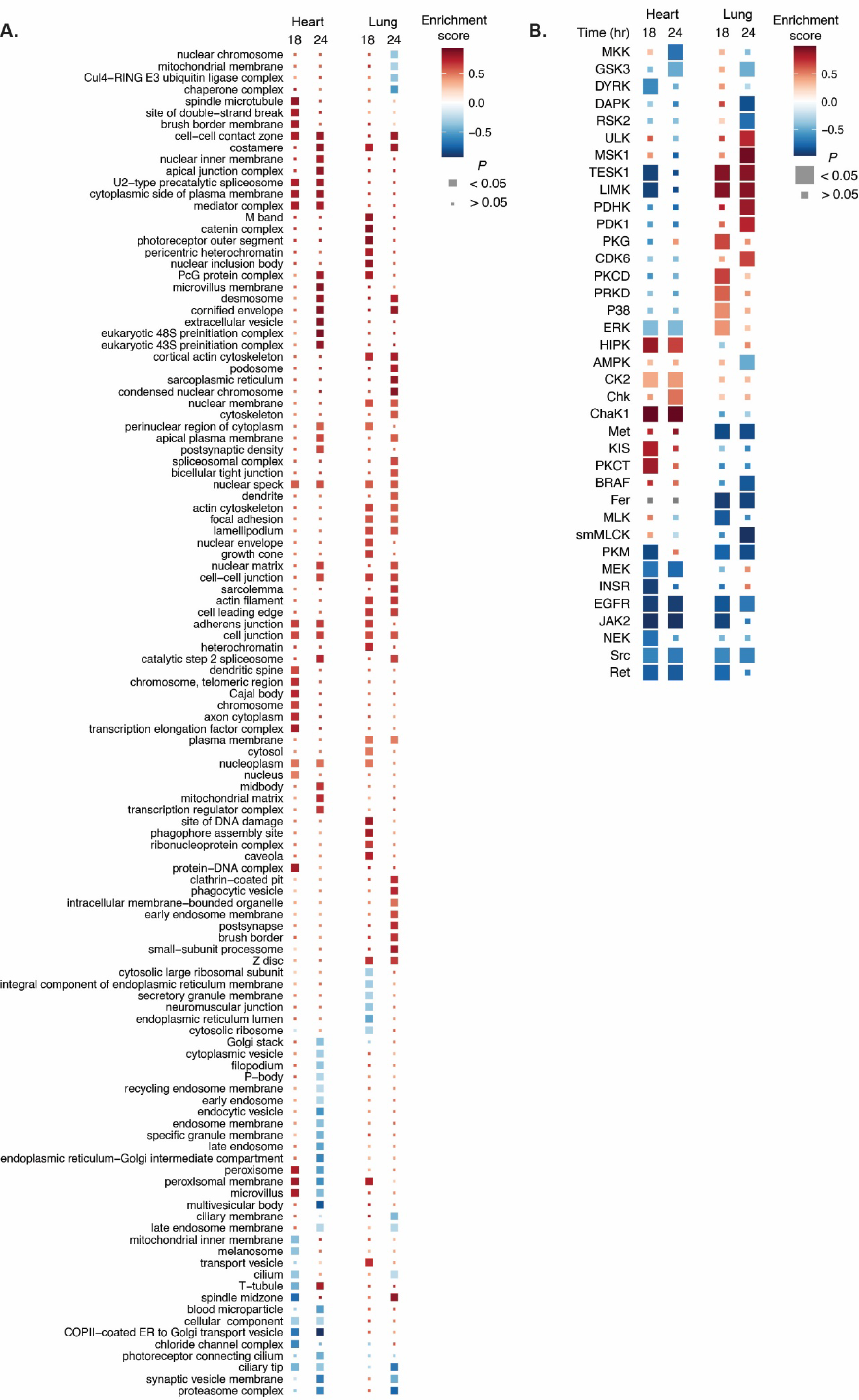
(A) All GO cellular components enriched in at least one condition (P < 0.05). (B) All kinases with reported substrates enriched in at least one condition (P < 0.05).

**Figure S7.**
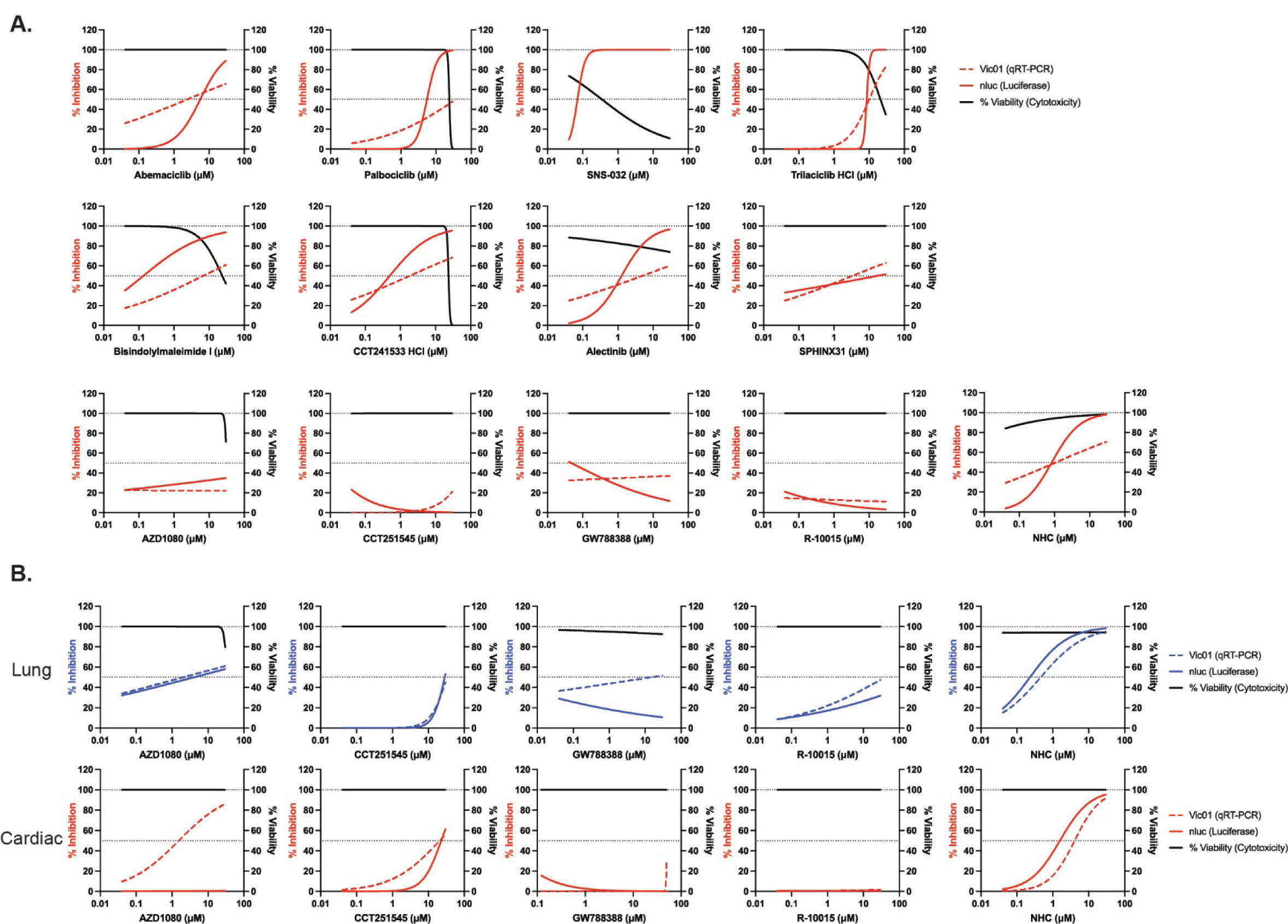
Cell viability (% relative to Vehicle control, black lines) and inhibition of SARS- CoV-2 growth (% relative to Vehicle control) in the presence of indicated compounds in H9 cardiac cells (A), H9 lung AT2 cells (B) and NKX2-5 cardiac (C) cells at 3 dpi.

**Table S1.** List of sites phosphorylated on SARS-CoV-2 proteins in infected cardiac and lung cells

| **Protein** | **Phosphorylation Site** | **Lung** | **Cardiac** |
| --- | --- | --- | --- |
| ORF1a | S2644 | + | + |
| Spike | S1261 | + | + |
| Membrane | S213 | + | + |
| Membrane | S214 | + | + |
| Membrane | T208 | - | + |
| Membrane | S212 | - | + |
| Nucleocapsid | S2 | + | + |
| Nucleocapsid | S21 | + | + |
| Nucleocapsid | S23 | + | + |
| Nucleocapsid | T24 | + | + |
| Nucleocapsid | S26 | + | + |
| Nucleocapsid | T76 | - | - |
| Nucleocapsid | T79 | + | + |
| Nucleocapsid | T166 | + | + |
| Nucleocapsid | Y172 | + | - |
| Nucleocapsid | S176 | + | + |
| Nucleocapsid | S180 | + | + |
| Nucleocapsid | S183 | - | - |
| Nucleocapsid | S184 | + | + |
| Nucleocapsid | S186 | - | - |
| Nucleocapsid | S188 | - | - |
| Nucleocapsid | S194 | + | + |
| Nucleocapsid | S197 | + | + |
| Nucleocapsid | T198 | + | + |
| Nucleocapsid | S201 | + | + |
| Nucleocapsid | S202 | + | + |
| Nucleocapsid | T205 | + | + |
| Nucleocapsid | S206 | + | + |
| Nucleocapsid | T245 | + | + |
| Nucleocapsid | T379 | + | + |
| Nucleocapsid | T391 | + | + |
| Nucleocapsid | T393 | + | + |
| Nucleocapsid | S404 | + | + |
| Nucleocapsid | S412 | + | + |
| Nucleocapsid | S416 | + | + |
| ORF9b | S50 | + | + |

**Table S2.** Compounds for antiviral testing from MedChemExpress

| **Compound** | **Target Kinase(s)** | **Catalogue** |
| --- | --- | --- |
| Abemaciclib | CDK4 and CDK6 | HY-16297A |
| Alectinib (CH5424802) | SRPK1 | HY-13011 |
| AZD1080 | GSK-3 | HY-13862 |
| Bisindolylmaleimide I | PKC | HY-13867 |
| CCT241533 hydrochloride | CHK2 | HY-14715B |
| CCT251545 | CDK8, CDK19 and WNT | HY-12681 |
| Dinaciclib | CDK2, CDK5, CDK1, and CDK9 | HY-10492 |
| Flavopiradol | CDK1, CDK2, CDK4 | HY-10005 |
| GW788388 | ALK5 and TGF-β type II receptor | HY-10326 |
| Palbociclib hydrochloride | CDK and CDK6 | HY-50767A |
| R-10015 | LIMK | HY-120097 |
| R406 free base | Syk/FLT3 | HY-11108 |
| Samuraciclib hydrochloride hydrate | CDK7 | HY-103712B |
| SNS-032 | CDK2, CDK7, and CDK9 | HY-10008 |
| SPHINX31 | SRPK1 | HY-117661 |
| Trilaciclib hydrochloride | CDK4 and CDK6 | HY-101467A |

ALK5: Activin-like kinase 5, CDK: Cyclin dependent kinase, CHK: Checkpoint, FLT3: FMS-like tyrosine kinase 3, GSK-3: Glycogen synthase kinase 3, LIMK: LIMK domain kinase, PKC: Protein kinase C, SRPK1: Serine/arginine-rich protein-specific kinase, TGF- $\beta$ : Transforming growth factor beta.

**Table S3.** TaqMan Probes for qPCR

| **Gene** | **Target Kinase(s)** |
| --- | --- |
| GAPDH | Hs02758991_g1, Hs02786624_g1 |
| TMPRSS2 | Hs00237175_m1 |
| ACE2 | Hs01085333_m1 |
| CTSL | Hs00964650_m1 |
| EIF4E | Hs00854166_g1 |
| IFN-β1 | Hs01077958_s1 |
| IL-6 | Hs00174131_m1 |
| IRF9 | Hs00196051_m1 |
| IRS1 | Hs00178563_m1 |
| STAT1 | Hs01013996_m1 |
| STAT1 | Hs01013116_g1 |
| IRS2 | Hs00275843_s1 |
| PRKCD | Hs01090047_m1 |
| SOCS1 | Hs00705164_s1 |
| REL | Hs00968440_m1 |
| MAP2K6 | Hs00992389_m1 |
| IFITM3 | Hs03057129_s1 |
| OAS2 | Hs00942643_m1 |

## References

1. Chen PZ, Bobrovitz N, Premji ZA, Koopmans M, Fisman DN, and Gu FX. SARS-CoV-2 shedding dynamics across the respiratory tract, sex, and disease severity for adult and pediatric COVID-19. Elife. 2021;10.

2. Hou YJ, Okuda K, Edwards CE, Martinez DR, Asakura T, Dinnon KH, 3rd, et al. SARS-CoV-2 Reverse Genetics Reveals a Variable Infection Gradient in the Respiratory Tract. Cell. 2020;182(2):429–46 e14.

3. Huang C, Huang L, Wang Y, Li X, Ren L, Gu X, et al. 6-month consequences of COVID-19 in patients discharged from hospital: a cohort study. Lancet. 2021;397(10270):220–32.

4. Nalbandian A, Sehgal K, Gupta A, Madhavan MV, McGroder C, Stevens JS, et al. Post-acute COVID-19 syndrome. Nat Med. 2021;27(4):601–15.

5. Lazzerini PE, Laghi-Pasini F, Boutjdir M, and Capecchi PL. Inflammatory cytokines and cardiac arrhythmias: the lesson from COVID-19. Nat Rev Immunol. 2022;22(5):270–2.

6. Shi S, Qin M, Shen B, Cai Y, Liu T, Yang F, et al. Association of Cardiac Injury With Mortality in Hospitalized Patients With COVID-19 in Wuhan, China. JAMA Cardiol. 2020;5(7):802–10.

7. Puntmann VO, Carerj ML, Wieters I, Fahim M, Arendt C, Hoffmann J, et al. Outcomes of Cardiovascular Magnetic Resonance Imaging in Patients Recently Recovered From Coronavirus Disease 2019 (COVID-19). JAMA Cardiol. 2020;5(11):1265–73.

8. Goyal P, Choi JJ, Pinheiro LC, Schenck EJ, Chen R, Jabri A, et al. Clinical Characteristics of Covid-19 in New York City. N Engl J Med. 2020;382(24):2372–4.

9. Shao MJ, Shang LX, Luo JY, Shi J, Zhao Y, Li XM, et al. Myocardial injury is associated with higher mortality in patients with coronavirus disease 2019: a meta- analysis. J Geriatr Cardiol. 2020;17(4):224–8.

10. Deinhardt-Emmer S, Wittschieber D, Sanft J, Kleemann S, Elschner S, Haupt KF, et al. Early postmortem mapping of SARS-CoV-2 RNA in patients with COVID-19 and the correlation with tissue damage. Elife. 2021;10.

11. Wang XM, Mannan R, Xiao L, Abdulfatah E, Qiao Y, Farver C, et al. Characterization of SARS-CoV-2 and host entry factors distribution in a COVID-19 autopsy series. Commun Med (Lond). 2021;1:24.

12. Schneider J, Pease D, Navaratnarajah C, Halfmann P, Clemens D, Ye D, et al. SARS-CoV-2 direct cardiac damage through spike-mediated cardiomyocyte fusion. Research Square. 2020.

13. Zhou P, Yang XL, Wang XG, Hu B, Zhang L, Zhang W, et al. A pneumonia outbreak associated with a new coronavirus of probable bat origin. Nature. 2020;579(7798):270–3.

14. Shang J, Wan Y, Luo C, Ye G, Geng Q, Auerbach A, et al. Cell entry mechanisms of SARS-CoV-2. Proc Natl Acad Sci U S A. 2020;117(21):11727–34.

15. Hoffmann M, Kleine-Weber H, Schroeder S, Kruger N, Herrler T, Erichsen S, et al. SARS-CoV-2 Cell Entry Depends on ACE2 and TMPRSS2 and Is Blocked by a Clinically Proven Protease Inhibitor. Cell. 2020;181(2):271–80 e8.

16. Ou X, Liu Y, Lei X, Li P, Mi D, Ren L, et al. Characterization of spike glycoprotein of SARS-CoV-2 on virus entry and its immune cross-reactivity with SARS-CoV. Nat Commun. 2020;11(1):1620.

17. Perez-Bermejo JA, Kang S, Rockwood SJ, Simoneau CR, Joy DA, Silva AC, et al. SARS-CoV-2 infection of human iPSC-derived cardiac cells reflects cytopathic features in hearts of patients with COVID-19. Sci Transl Med. 2021;13(590).

18. Sungnak W, Huang N, Becavin C, Berg M, Queen R, Litvinukova M, et al. SARS-CoV-2 entry factors are highly expressed in nasal epithelial cells together with innate immune genes. Nat Med. 2020;26(5):681–7.

19. Qi F, Qian S, Zhang S, and Zhang Z. Single cell RNA sequencing of 13 human tissues identify cell types and receptors of human coronaviruses. Biochem Biophys Res Commun. 2020;526(1):135–40.

20. Zou X, Chen K, Zou J, Han P, Hao J, and Han Z. Single-cell RNA-seq data analysis on the receptor ACE2 expression reveals the potential risk of different human organs vulnerable to 2019-nCoV infection. Front Med. 2020;14(2):185–92.

21. Hamming I, Timens W, Bulthuis ML, Lely AT, Navis G, and van Goor H. Tissue distribution of ACE2 protein, the functional receptor for SARS coronavirus. A first step in understanding SARS pathogenesis. J Pathol. 2004;203(2):631–7.

22. Muus C, Luecken MD, Eraslan G, Sikkema L, Waghray A, Heimberg G, et al. Single-cell meta-analysis of SARS-CoV-2 entry genes across tissues and demographics. Nat Med. 2021;27(3):546–59.

23. Chen J, Neil J, Tan J, Rudraraju R, Mohenska M, Sun Y, et al. An iTSC-derived placental model of SARS-CoV-2 infection reveals ACE2-dependent susceptibility in syncytiotrophoblasts. bioRxiv. 2021:2021.10.27.465224.

24. Huang J, Hume AJ, Abo KM, Werder RB, Villacorta-Martin C, Alysandratos KD, et al. SARS-CoV-2 Infection of Pluripotent Stem Cell-Derived Human Lung Alveolar Type 2 Cells Elicits a Rapid Epithelial-Intrinsic Inflammatory Response. Cell Stem Cell. 2020;27(6):962–73 e7.

25. Sharma A, Garcia G, Jr., Wang Y, Plummer JT, Morizono K, Arumugaswami V, et al. Human iPSC-Derived Cardiomyocytes Are Susceptible to SARS-CoV-2 Infection. Cell Rep Med. 2020;1(4):100052.

26. Williams TL, Colzani MT, Macrae RGC, Robinson EL, Bloor S, Greenwood EJD, et al. Human embryonic stem cell-derived cardiomyocyte platform screens inhibitors of SARS-CoV-2 infection. Commun Biol. 2021;4(1):926.

27. Anderson DJ, Kaplan DI, Bell KM, Koutsis K, Haynes JM, Mills RJ, et al. NKX2-5 regulates human cardiomyogenesis via a HEY2 dependent transcriptional network. Nat Commun. 2018;9(1):1373.

28. Lopes LR, Garcia-Hernandez S, Lorenzini M, Futema M, Chumakova O, Zateyshchikov D, et al. Alpha-protein kinase 3 (ALPK3) truncating variants are a cause of autosomal dominant hypertrophic cardiomyopathy. Eur Heart J. 2021;42(32):3063–73.

29. Jacob A, Morley M, Hawkins F, McCauley KB, Jean JC, Heins H, et al. Differentiation of Human Pluripotent Stem Cells into Functional Lung Alveolar Epithelial Cells. Cell Stem Cell. 2017;21(4):472–88 e10.

30. Elliott DA, Braam SR, Koutsis K, Ng ES, Jenny R, Lagerqvist EL, et al. NKX2- 5(eGFP/w) hESCs for isolation of human cardiac progenitors and cardiomyocytes. Nat Methods. 2011;8(12):1037–40.

31. Phelan DG, Anderson DJ, Howden SE, Wong RC, Hickey PF, Pope K, et al. ALPK3-deficient cardiomyocytes generated from patient-derived induced pluripotent stem cells and mutant human embryonic stem cells display abnormal calcium handling and establish that ALPK3 deficiency underlies familial cardiomyopathy. Eur Heart J. 2016;37(33):2586–90.

32. Humphrey SJ, Karayel O, James DE, and Mann M. High-throughput and high- sensitivity phosphoproteomics with the EasyPhos platform. Nat Protoc. 2018;13(9):1897–916.

33. Yaron TM, Heaton BE, Levy TM, Johnson JL, Jordan TX, Cohen BM, et al. The FDA-approved drug Alectinib compromises SARS-CoV-2 nucleocapsid phosphorylation and inhibits viral infection in vitro. bioRxiv. 2020.

34. Zhou P, Yang XL, Wang XG, Hu B, Zhang L, Zhang W, et al. Addendum: A pneumonia outbreak associated with a new coronavirus of probable bat origin. Nature. 2020;588(7836):E6.

35. Ziegler CGK, Allon SJ, Nyquist SK, Mbano IM, Miao VN, Tzouanas CN, et al. SARS-CoV-2 Receptor ACE2 Is an Interferon-Stimulated Gene in Human Airway Epithelial Cells and Is Detected in Specific Cell Subsets across Tissues. Cell. 2020;181(5):1016–35 e19.

36. Marchiano S, Hsiang TY, Khanna A, Higashi T, Whitmore LS, Bargehr J, et al. SARS-CoV-2 Infects Human Pluripotent Stem Cell-Derived Cardiomyocytes, Impairing Electrical and Mechanical Function. Stem Cell Reports. 2021;16(3):478–92.

37. Bojkova D, Wagner JUG, Shumliakivska M, Aslan GS, Saleem U, Hansen A, et al. SARS-CoV-2 infects and induces cytotoxic effects in human cardiomyocytes. Cardiovasc Res. 2020;116(14):2207–15.

38. Bailey AL, Dmytrenko O, Greenberg L, Bredemeyer AL, Ma P, Liu J, et al. SARS- CoV-2 Infects Human Engineered Heart Tissues and Models COVID-19 Myocarditis. JACC Basic Transl Sci. 2021;6(4):331–45.

39. Montaser M, Lalmanach G, and Mach L. CA-074, but not its methyl ester CA- 074Me, is a selective inhibitor of cathepsin B within living cells. Biol Chem. 2002;383(7-8):1305–8.

40. Tiwari SK, Wang S, Smith D, Carlin AF, and Rana TM. Revealing Tissue-Specific SARS-CoV-2 Infection and Host Responses using Human Stem Cell-Derived Lung and Cerebral Organoids. Stem Cell Reports. 2021;16(3):437–45.

41. Gunst JD, Staerke NB, Pahus MH, Kristensen LH, Bodilsen J, Lohse N, et al. Efficacy of the TMPRSS2 inhibitor camostat mesilate in patients hospitalized with Covid-19-a double-blind randomized controlled trial. EClinicalMedicine. 2021;35:100849.

42. Ou T, Mou H, Zhang L, Ojha A, Choe H, and Farzan M. Hydroxychloroquine- mediated inhibition of SARS-CoV-2 entry is attenuated by TMPRSS2. PLoS Pathog. 2021;17(1):e1009212.

43. Mills RJ, Titmarsh DM, Koenig X, Parker BL, Ryall JG, Quaife-Ryan GA, et al. Functional screening in human cardiac organoids reveals a metabolic mechanism For cardiomyocyte cell cycle arrest. Proc Natl Acad Sci USA. 2017;114(40):E8372–E81.

44. Li Y, Renner DM, Comar CE, Whelan JN, Reyes HM, Cardenas-Diaz FL, et al. SARS-CoV-2 induces double-stranded RNA-mediated innate immune responses in respiratory epithelial-derived cells and cardiomyocytes. Proc Natl Acad Sci USA. 2021;118(16).

45. Sampaio NG, Chauveau L, Hertzog J, Bridgeman A, Fowler G, Moonen JP, et al. The RNA sensor MDA5 detects SARS-CoV-2 infection. Sci Rep. 2021;11(1):13638.

46. Shemesh M, Aktepe TE, Deerain JM, McAuley JL, Audsley MD, David CT, et al. SARS-CoV-2 suppresses IFNbeta production mediated by NSP1, 5, 6, 15, ORF6 and ORF7b but does not suppress the effects of added interferon. PLoS Pathog. 2021;17(8):e1009800.

47. Lei X, Dong X, Ma R, Wang W, Xiao X, Tian Z, et al. Activation and evasion of type I interferon responses by SARS-CoV-2. Nat Commun. 2020;11(1):3810.

48. Xia H, Cao Z, Xie X, Zhang X, Chen JY, Wang H, et al. Evasion of Type I Interferon by SARS-CoV-2. Cell Rep. 2020;33(1):108234.

49. Uhlen M, Fagerberg L, Hallstrom BM, Lindskog C, Oksvold P, Mardinoglu A, et al. Proteomics. Tissue-based map of the human proteome. Science. 2015;347(6220):1260419.

50. Zhang Q, Bastard P, Liu Z, Le Pen J, Moncada-Velez M, Chen J, et al. Inborn errors of type I IFN immunity in patients with life-threatening COVID-19. Science. 2020;370(6515).

51. Pruijssers AJ, George AS, Schafer A, Leist SR, Gralinksi LE, Dinnon KH, 3rd, et al. Remdesivir Inhibits SARS-CoV-2 in Human Lung Cells and Chimeric SARS- CoV Expressing the SARS-CoV-2 RNA Polymerase in Mice. Cell Rep. 2020;32(3):107940.

52. Bouhaddou M, Memon D, Meyer B, White KM, Rezelj VV, Correa Marrero M, et al. The Global Phosphorylation Landscape of SARS-CoV-2 Infection. Cell. 2020;182(3):685–712 e19.

53. Tohyama S, Hattori F, Sano M, Hishiki T, Nagahata Y, Matsuura T, et al. Distinct metabolic flow enables large-scale purification of mouse and human pluripotent stem cell-derived cardiomyocytes. Cell Stem Cell. 2013;12(1):127–37.

54. Howe KL, Achuthan P, Allen J, Allen J, Alvarez-Jarreta J, Amode MR, et al. Ensembl 2021. Nucleic Acids Res. 2021;49(D1):D884–D91.

55. Dobin A, Davis CA, Schlesinger F, Drenkow J, Zaleski C, Jha S, et al. STAR: ultrafast universal RNA-seq aligner. Bioinformatics. 2013;29(1):15–21.

56. Liao Y, Smyth GK, and Shi W. featureCounts: an efficient general purpose program for assigning sequence reads to genomic features. Bioinformatics. 2014;30(7):923–30.

57. Robinson MD, McCarthy DJ, and Smyth GK. edgeR: a Bioconductor package for differential expression analysis of digital gene expression data. Bioinformatics. 2010;26(1):139–40.

58. Zhou Y, Zhou B, Pache L, Chang M, Khodabakhshi AH, Tanaseichuk O, et al. Metascape provides a biologist-oriented resource for the analysis of systems-level datasets. Nat Commun. 2019;10(1):1523.

